# Tissue Tropism and Transmission Ecology Predict Virulence of Human RNA Viruses

**DOI:** 10.1101/581512

**Authors:** Liam Brierley, Amy B. Pedersen, Mark E. J. Woolhouse

**Author notes:** **Corresponding author:**, Current address: sigma, Coventry University, Priory Street, Coventry, CV1 5FB, UK.

## Abstract

Novel infectious diseases continue to emerge within human populations. Predictive studies have begun to identify pathogen traits associated with emergence. However, emerging pathogens vary widely in virulence, a key determinant of their ultimate risk to public health. Here, we use structured literature searches to review the virulence of each of the 214 known human-infective RNA virus species. We then use a machine learning framework to determine whether viral virulence can be predicted by ecological traits including human-to-human transmissibility, transmission routes, tissue tropisms and host range. Using severity of clinical disease as a measurement of virulence, we identified potential risk factors using predictive classification tree and random forest ensemble models. The random forest model predicted literature-assigned disease severity of test data with 90.3% accuracy, compared to a null accuracy of 74.2%. In addition to viral taxonomy, the ability to cause systemic infection, having renal and/or neural tropism, direct contact or respiratory transmission, and limited (0 < R_0_ ≤ 1) human-to-human transmissibility were the strongest predictors of severe disease. We present a novel, comparative perspective on the virulence of all currently known human RNA virus species. The risk factors identified may provide novel perspectives in understanding the evolution of virulence and elucidating molecular virulence mechanisms. These risk factors could also improve planning and preparedness in public health strategies as part of a predictive framework for novel human infections.

**Author Summary:** Newly emerging infectious diseases present potentially serious threats to global health. Although studies have begun to identify pathogen traits associated with the emergence of new human diseases, these do not address why emerging infections vary in the severity of disease they cause, often termed ‘virulence’. We test whether ecological traits of human viruses can act as predictors of virulence, as suggested by theoretical studies. We conduct the first systematic review of virulence across all currently known human RNA virus species. We adopt a machine learning approach by constructing a random forest, a model that aims to optimally predict an outcome using a specific structure of predictors. Predictions matched literature-assigned ratings for 28 of 31 test set viruses. Our predictive model suggests that higher virulence is associated with infection of multiple organ systems, nervous systems or the renal systems. Higher virulence was also associated with contact-based or airborne transmission, and limited capability to transmit between humans. These risk factors may provide novel starting points for questioning why virulence should evolve and identifying causative mechanisms of virulence. In addition, our work could suggest priority targets for infectious disease surveillance and future public health risk strategies.

**Blurb:** Comparative analysis using machine learning shows specificity of tissue tropism and transmission biology can act as predictive risk factors for virulence of human RNA viruses.

## Introduction

The emergence of novel infectious diseases continues to represent a threat to global public health. Emerging pathogens have been defined as those newly recognised infections of humans following zoonotic transmission, or those increasing in incidence and/or geographic range [1]. High-profile examples of emerging pathogens include the discovery of the novel MERS coronavirus from cases of respiratory illness in 2012 [2], and the expansion of the range of Zika virus across the South Pacific and the Americas [3]. The emergence of previously unseen viruses means that the set of known human viruses continually increases by around 2 species per year [4,5]. Initial comparative studies identified trends among emerging human pathogens, for example, increased risk of emergence for pathogens with broad host ranges, and RNA viruses [6–9]. However, more recent comparative analyses have focused on risk factors for specific pathogen traits, such as transmissibility [10–12]. Here, we focus on understanding the ecological determinants of pathogen virulence, using all currently recognised human RNA viruses as a study system.

Emerging RNA viruses vary widely in their virulence, with some never having been associated with human disease at all. For example, Zaire ebolavirus causes severe haemorrhagic fever with outbreaks, including the 2014 West African outbreak showing case fatality ratios of ∼60% or more [13,14]. In contrast, human infections with Reston ebolavirus have never exhibited any evidence of disease symptoms [15]. Applying the comparative approach to understand the ecology of virulence could offer valuable synergy with studies of emergence, towards prioritisation and preparedness in the detection of potential new human viruses [16].

Few comparative analyses have addressed the risk factors driving human pathogen virulence to date (but see [17–19]), and none have exhaustively investigated virulence across the breadth of all currently recognised human RNA viruses. Several hypotheses regarding how pathogen ecology affects virulence have been derived from theoretical models of evolution. For example, the trade-off hypothesis was developed based on the assumption that rate of transmission between individuals may increase as a function of virulence, but there will be a consequential increase in host mortality (or decrease in host recovery as the inverse of mortality). As a result, pathogen fitness will be subject to trade-off between virulence and transmissibility over a longer infectious window [20,21]. The trade-off hypothesis is highly debated as it is difficult to empirically characterise due to dependency on many other aspects of host-pathogen coevolution [22,23]. However, comparative analysis has been suggested as one method to assess evidence for a virulence-transmission trade-off [22]. Based on these core principles, we hypothesised that limited capability to transmit between humans may act as a predictive risk factor for virulence. We also note that evolutionary trade-offs will only apply to coevolved host-virus relationships and that many human viruses result from zoonotic cross-species transmission without onward transmission or adaptation. In these cases, ‘coincidental’ non-adapted virulence may result [24,25], and as above, we hypothesised that limited human-to-human transmissibility may predict higher virulence.

Transmission route may also influence the evolution of virulence. Ewald [18] suggested that vector-borne pathogens should be less constrained by costs of virulence, i.e. morbidity and immobilisation of the vertebrate host does not impede transmission if it occurs through an arthropod vector. We therefore hypothesised a vector-borne transmission route would predict higher virulence.

Several studies have also suggested a link between host range and virulence. Assuming an evolutionary trade-off exists between virulence and transmission rate, higher virulence may result in pathogens with narrower host ranges following selection pressures to increase transmission rate within the specialist host(s) [19]. Furthermore, the degree of virulence in experimental infections with *Drosophila C virus* was more similar between closely related hosts [26]. Though similar ideas have not yet been formally tested for human infections, parasite infectivity correlates with phylogenetic relatedness among primates [27]. We hypothesised infection of non-human primates as a specific related host taxon would predict higher virulence. Finally, although yet unexplored via theoretical models, it may be an intuitive expectation that systemic infections present with more severe disease than local infections. A broader tissue tropism could therefore also predict higher virulence.

We aimed to determine patterns of virulence across the breadth of all known human RNA viruses. We then aimed to use predictive machine learning models to ask whether ecological traits of viruses can act as predictive risk factors for virulence in humans. Specifically, we examined hypotheses that viruses would be more highly virulent if they: lacked transmissibility within humans; had vector-borne transmission routes; had a narrow host range including non-human primates; or had greater breadth of tissue tropisms.

## Results

### Virulence of Human RNA Viruses

Following [5], as of 2015 there were 214 RNA virus species containing viruses capable of infecting humans, spanning 55 genera and 21 families (with one species unassigned to a family). Using a two-category system, 58 of these were rated as causing ‘severe’ clinical disease and 154 as ‘nonsevere’ following systematic literature review (Fig 2, see also S1 Table, S2 Table). Two virus species could not be assigned a disease severity rating and were excluded from all analyses (*Hepatitis delta virus*, which is reliant on *Hepatitis B virus* coinfection; and *Primate T-lymphotropic virus 3*, which may be associated with chronic disease like other T-lymphotropic viruses, but has not been known in humans long enough for cohort observations). Disease severity differed between viral taxonomic families (Fisher’s exact, 1000 simulations, p < 0.001), with *Arenaviridae*, *Filoviridae* and *Hantaviridae* having the highest fractions of severe-rated virus species (Fig 2). Fatalities were reported in healthy adults for 64 viruses and in vulnerable individuals only for an additional 26 viruses, whilst 8 viruses rated ‘nonsevere’ had severe strains, 6 of which belonged to the family *Picornaviridae*.

**Fig 1.**
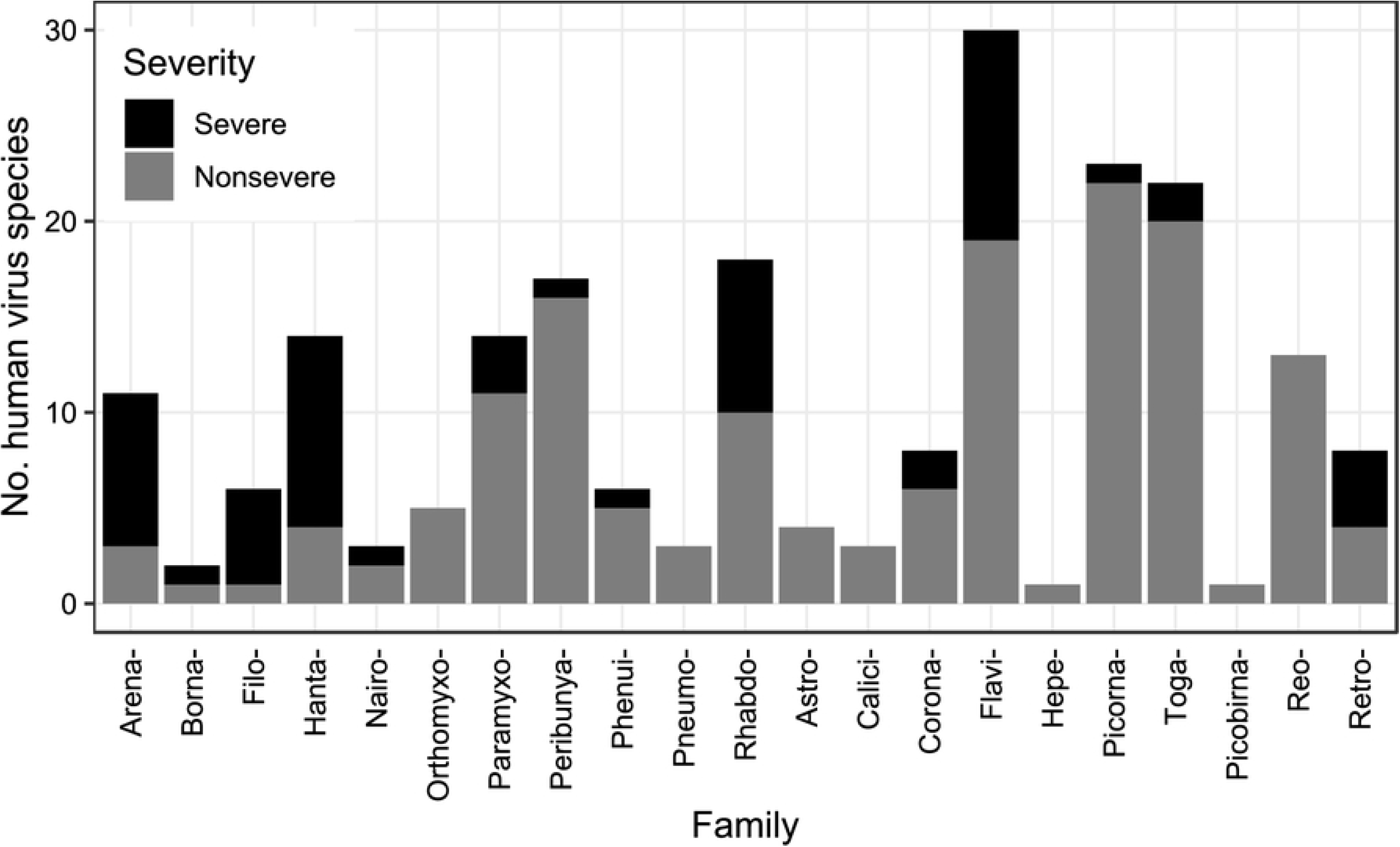
Virulence of currently known human RNA viruses with respect to taxonomy. Number of known human RNA virus species split by ICTV taxonomic family. Shading denotes disease severity rating.

**Fig 2.**
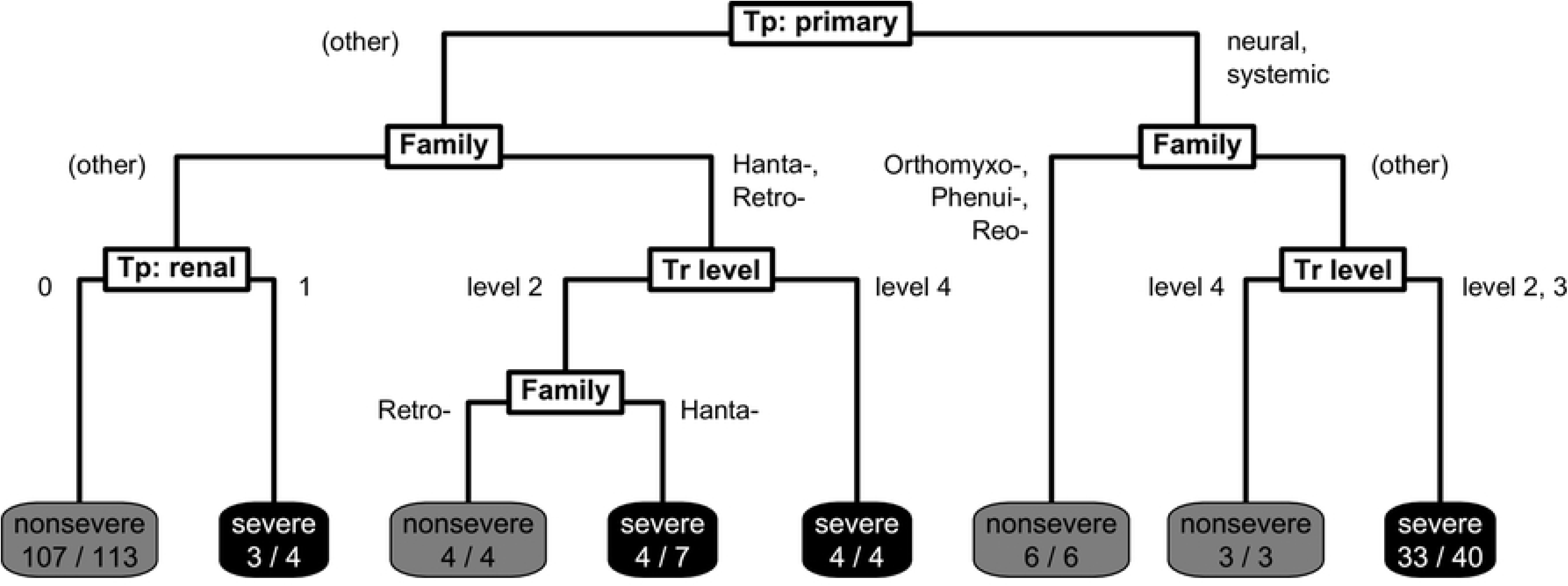
Final pruned classification tree predicting disease severity for 181 human RNA viruses. Final classification tree structure predicting virulence. Viruses begin at the top and are classified according to split criteria (white boxes) until reaching terminal nodes with the model’s prediction of disease severity, and the fraction of viruses following that path correctly classified, based on literature-assigned ratings (shaded boxes). ‘Tp: primary’ denotes primary tissue tropism, ‘Tr level’ denotes level of human-to-human transmissibility, and ‘Tp: renal.’ denotes having a known renal tissue tropism.

### Classification Tree Risk Factor Analysis

To find predictive risk factors for virulence, we firstly divided the 212 virus species into a training set (n = 181) and test set (n = 31) based on taxonomy and severity in order to minimise potential biases from trait imbalances. Using the training set, we then constructed a single classification tree that aimed to optimally classify viruses in virulence based on their ecological traits. The final pruned classification tree included variables relating to transmissibility, tissue tropism and taxonomy (Fig 2). Severe disease was predicted by the model for four generalised groups: i) viruses with a neural or systemic primary tropism with limited human-to-human transmissibility (excluding orthomyxoviruses, phenuiviruses and reoviruses); ii) viruses known to have a renal tropism (primary or otherwise); iii) hantaviruses; and iv) retroviruses with sustained human-to-human transmissibility.

### Random Forest Risk Factor Analysis

Although the illustrated classification tree identified several risk factors, this represents one of many possible trees, as tree structure is dependent on the exact sampling partition between training and test data. We therefore constructed a random forest model containing 5000 individual trees, each built using a bootstrapped sample of the training data and a randomly restricted subset of predictors.

Aggregated over these bootstrapped trees, the most informative predictor variables for classifying virulence were taxonomic family and primary tissue tropism (Fig 4). However, transmission route, human-to-human transmissibility level, and having a known neural or renal tropism were also relatively informative, broadly mirroring the risk factors observed in the single tree. Host range predictors were generally uninformative.

**Fig 3.**
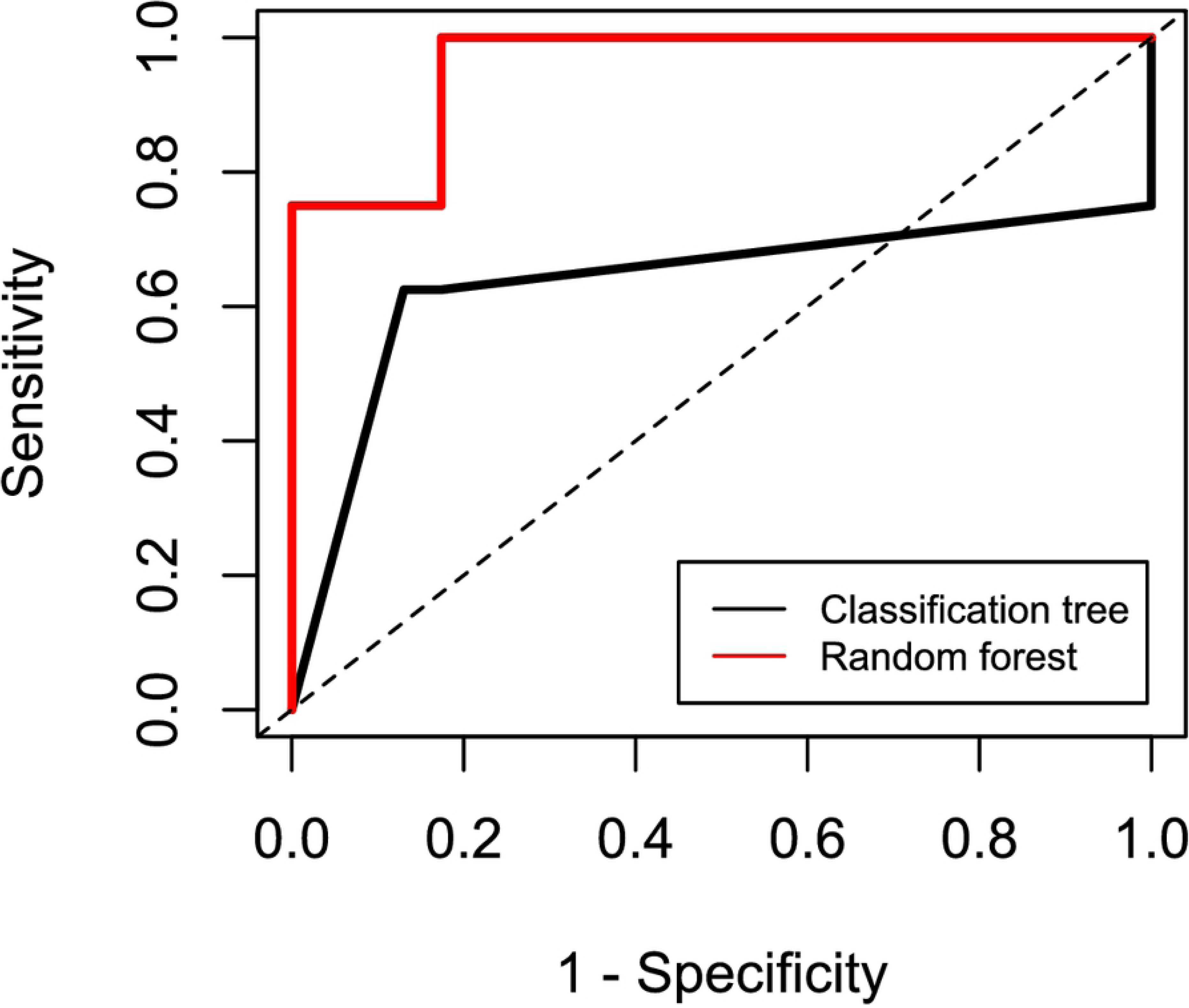
Receiver operating characteristic curve for tree-based machine learning models. Plotted model predictive performance for the single classification tree (bold black line) and the random forest (bold red line) models when applied to the test set. Y axis denotes sensitivity (or true positive rate; proportion of viruses rated ‘severe’ by literature protocol that were correctly predicted as ‘severe’ by the model), and X axis denotes 1 – specificity (or false positive rate; proportion of viruses rated ‘nonsevere’ by literature protocol that were incorrectly predicted as ‘severe’ by the model). Dashed black line indicates null expectation (i.e. a model with no discriminatory power). Model profiles further toward the top left indicate a better predictive performance.

**Fig 4.**
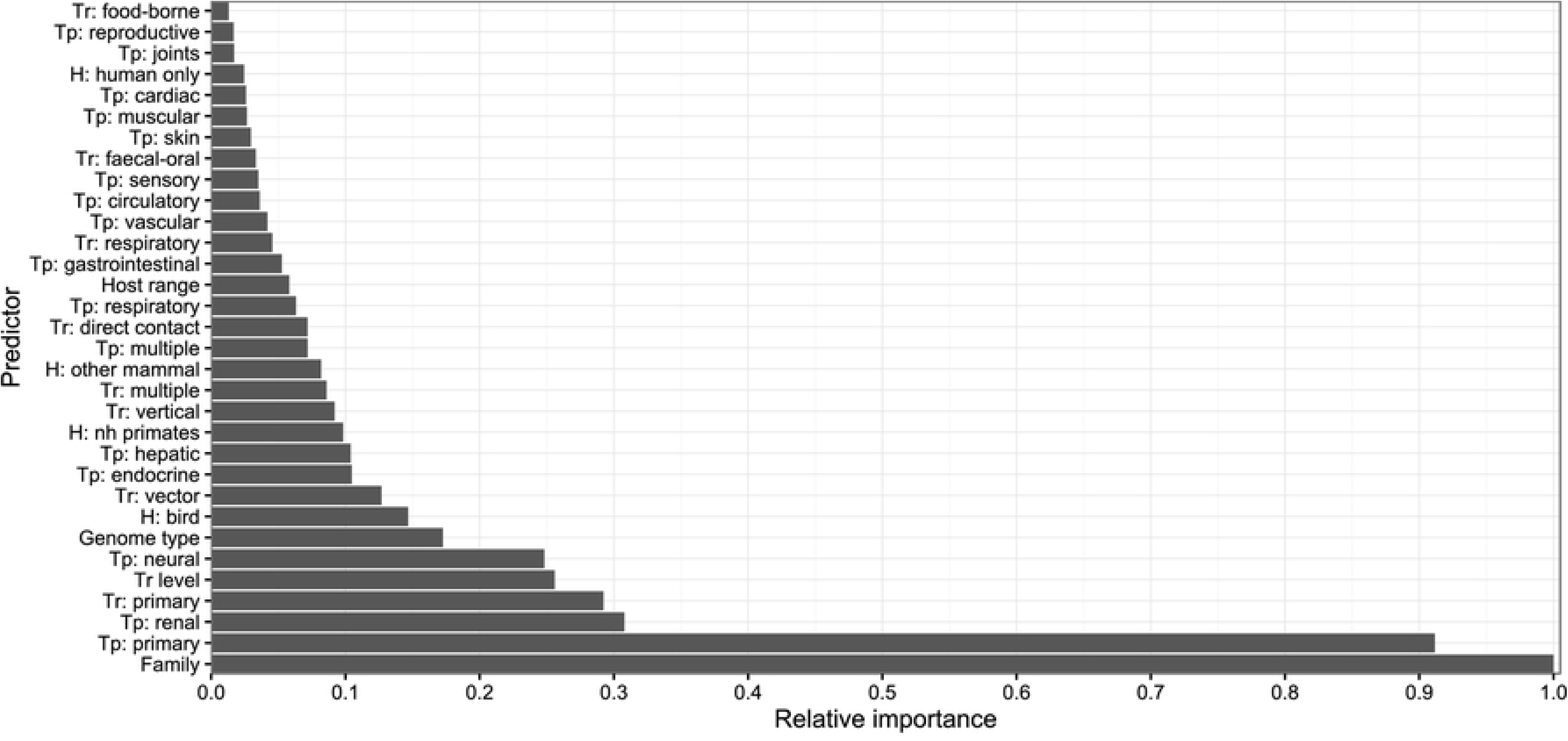
Variable importances from the random forest model. Importance of each predictor variable across the 5000 bootstrapped trees within the random forest, calculated as the mean decrease in Gini impurity following a tree split based on that predictor and scaled against the most informative predictor (taxonomic family) to give a relative measure. ‘Tp’ denotes tissue tropism predictor, ‘Tr’ denotes transmission route predictor, ‘Tr level’ denotes level of human-to-human transmissibility, and ‘H’ denotes host range predictor.

To quantify the effects of the most informative risk factors, partial dependences were extracted from the random forest, describing the marginal predicted probabilities of severe virulence associated with each virus trait (Fig 5, S3 Table). Averaging across other predictors, viruses having tissue tropisms within neural, renal or systemic across multiple organ systems presented the highest risk of severe virulence, whilst respiratory and gastrointestinal tropisms presented the lowest risk. An increased probability of severe virulence was also observed for viruses transmitted by direct contact or respiratory routes, and those with known but limited human-to-human transmissibility.

**Fig 5.**
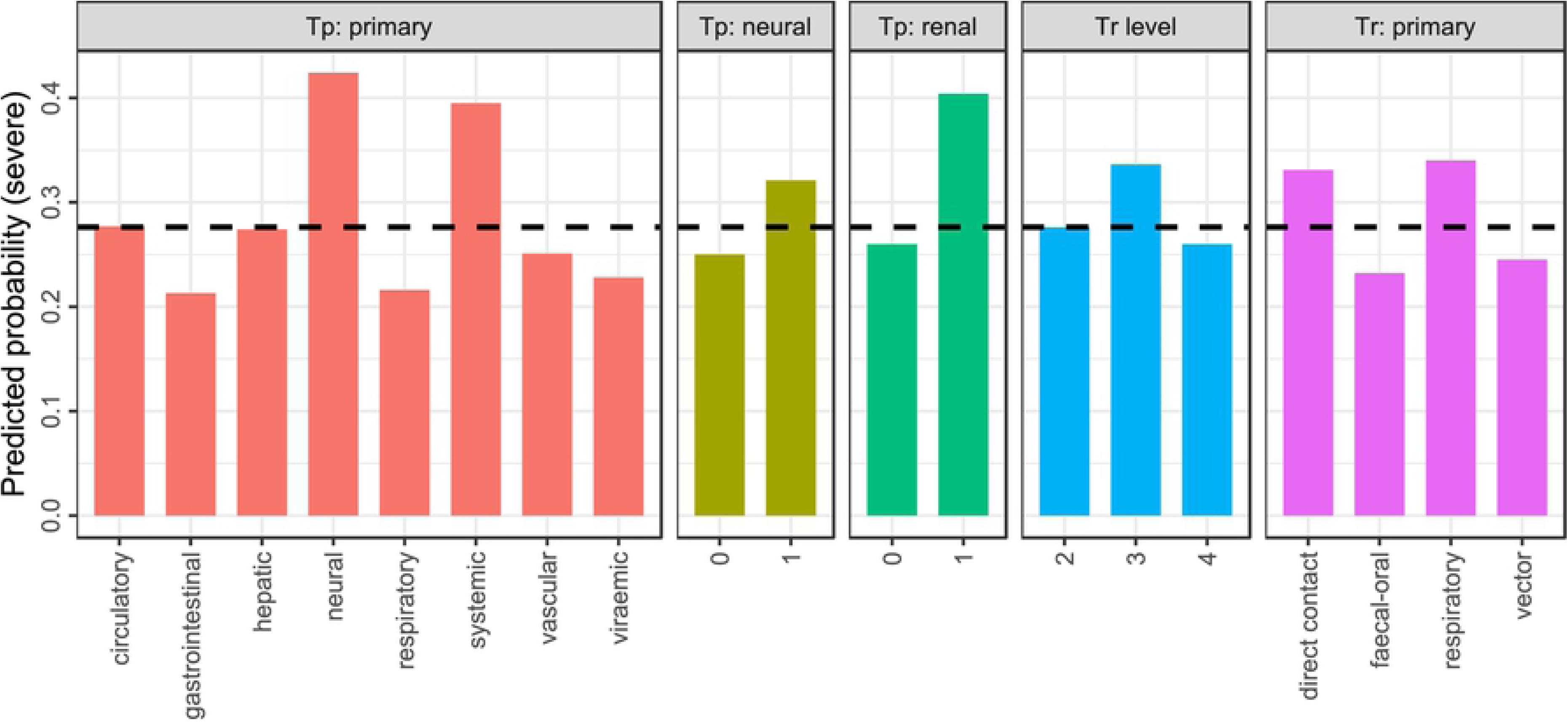
Partial dependences from the random forest model in predicting severe virulence. Predicted probability of classifying virulence as ‘severe’ for each of the most informative risk factors (primary tissue tropism, any known neural tropism, any known renal tropism, level of human-to-human transmissibility, and primary transmission route). Probabilities given are marginal, i.e. averaging over any effects of other predictors. Dashed line denotes raw prevalence of ‘severe’ virulence rating among the training dataset.

### Model Performance in Predicting Viral Virulence

Although the single classification tree model predicted the training set well, it did not appear generalisable to novel data within the test set. The single tree correctly predicted virulence ratings from literature-based criteria for 24 of 31 viruses in the test set giving a resulting accuracy of 77.4% (95% confidence interval [CI]: 58.9% - 90.4%), no evident improvement on the null model assigning all viruses as nonsevere (null accuracy = 74.2%). The random forest gave better predictive accuracy, correctly predicting virulence ratings for 28 of 31 test set viruses (accuracy: 90.3%, 95% CI: 74.3% - 98.0%), significantly greater than the null accuracy (exact binomial one-tailed test, p = 0.025). The random forest also achieved superior performance when considering sensitivity, specificity, True Skill Statistic, and the negative predictive value as a performance measure prioritising correct classification of ‘severe’-rated viruses (Table 1). The random forest also outperformed the classification tree in AUROC, area under the receiver operating characteristic curve (Table 1, Fig 3).

**Table 1.** Predictive performance metrics for classification tree and random forest model. Sensitivity, specificity, NPV (negative predictive value; proportion of ‘nonsevere’ predictions that correctly matched literature rating), TSS (true skill statistic; sensitivity + specificity – 1) and AUROC (area under receiver operating characteristic curve) for predictive model methods applied to predict virulence of 31 viruses within the test set.

| Model | Sensitivity | Specificity | NPV | TSS | AUROC |
| --- | --- | --- | --- | --- | --- |
| Classification tree | 0.625 | 0.826 | 0.864 | 0.451 | 0.636 |
| Random forest | 0.750 | 0.957 | 0.917 | 0.707 | 0.957 |

All misclassifications from the random forest occurred within the genus *Flavivirus* (S2 Table). Within the test set, there were two flaviruses rated as severe from literature protocols that were predicted to be nonsevere (*Rio Bravo virus, Yellow fever virus*), and one nonsevere flavivirus predicted to be severe (*Usutu virus*).

The observed predictor importances and risk factor directions were robust to constructing random forest models for subsets of viruses, removing those with low-certainty data or data from serological evidence only (S1 Fig, S2 Fig), and similar performance diagnostics were obtained (S5 Table). Redefining our virulence measure to integrate information on known fatalities and differences with subspecies or strains in an ordinal ranking system (S5 Table) did not improve predictive performance (S6 Table). Using alternative virulence measurements, the most informative variables and virus traits predicting severity showed good agreement with that of the main analysis (S3 Fig, S4 Fig) though when definitions of ‘severe’ virulence were widened, hepatic tropism became an informative predictor towards disease severity.

## Discussion

We present the first comparative analysis of virulence across all known human RNA virus species to our knowledge. We find that disease severity is non-randomly distributed across virus families and that beyond taxonomy, severe disease is predicted by risk factors of tissue tropism, and to a lesser extent, transmission route and level of human-to-human transmissibility. In both the classification tree and random forest, viruses were more likely to be predicted to cause severe disease if they caused systemic infections, had neural or renal tropism, transmitted via direct contact or respiratory routes, or had limited capability to transmit between humans (0 < R_0_ ≤ 1). These risk factors were robust to alternative modelling methods, alternative definitions of virulence, and exclusions of poor quality data.

### Ecology and Evolution of Risk Factor Traits

Primary tissue tropism was the most informative non-taxonomic risk factor (Fig 4) and the first split criteria in the classification tree (Fig 2), with specific neural tropism and generalised systemic tropism predicting severe disease (Fig 5). Few evolutionary studies have directly predicted how tissue tropism should influence virulence. The identified risk factor tropisms could be explainable as a simple function of pathology occurring in multiple or sensitive tissues respectively, increasing intensity of clinical disease. However, it has been suggested that an excessive, non-adapted virulence may result if infections occur within non-target tissues that do not contribute to transmission [28]. Furthermore, the evolutionary determinants of tissue tropism themselves are not well understood [29]. Tissue tropism should be a key consideration for future comparative and evolutionary modelling efforts.

We also found viruses primarily transmitted by direct contact and respiratory routes to have a higher predicted probability of severe virulence than viruses transmitted by more indirect faecal-oral or vector-borne routes. Contrastingly, Ewald [18] reported a positive association between virulence and vector-borne transmission in comparative analyses pooling several microparasite types, including a limited range of viruses, and suggested virulence has fewer costs to viral evolutionary fitness if vector transmission can occur independent of host health and mobility. The opposite association we observe may imply that even if transmission occurs via an indirect route such as through an arthropod vector, virulence could bring ultimate fitness costs due to host mortality before encountering a vector, fomite, etc..

The relationship between virulence and transmissibility appears more complex. Firstly, the random forest model suggested a lower risk of severe virulence for viruses with sustained human-to-human transmissibility (level 4) (Fig 5). This would lend support towards hypothesised virulence-transmissibility trade-offs [20–22] and suggests that the adaptation necessary to develop efficient human-to-human transmissibility could result in attenuation of virulence in RNA viruses. Sustained transmissibility appeared to positively predict severe disease for a specific subset of four viruses in the single classification tree (Fig 2), all retroviruses causing chronic syndromes (*HIV 1* and *2*, *Primate T-lymphotropic virus 1* and *2*), which are likely subject to different evolutionary dynamics – if disease occurs after the infectious period, virulence brings fewer costs to pathogens from host mortality, essentially ‘decoupling’ from transmission [24]. We note only three non-chronic level 4 viruses rated severe: *Severe acute respiratory syndrome-related coronavirus, Yellow fever virus*, and *Zaire ebolavirus*.

Secondly, cross-species infections incapable of onward transmission (sometimes termed ‘dead-end’ infections) have been predicted to result in higher virulence as without any evolutionary selection, viral phenotypes within that host will be non-adapted, i.e. a ‘coincidental’ by-product [24,25]. However, we did not observe viruses incapable of human-to-human transmissibility to be more virulent, the highest risk instead being observed for viruses with self-limited transmissibility. This may suggest that if virulence is entirely unselected in dead-end infections, ultimate levels of virulence could also feasibly turn out to be ‘coincidentally’ low.

Taxonomic family being a highly informative predictor in the random forest implies that there is a broad phylogenetic signal to virulence, but it is also highly likely that the explanatory power represents a proxy for many other phylogenetically-conserved viral traits that are challenging to implement in comparative analyses of this scale, such as variation at the proteomic, transcriptomic or genomic level; or further data beyond simple categorisations, e.g. specific arthropod vector species. Untangling these sources of variation from different scales of traits will be a critical next step in predictive modelling of viral virulence.

### Analytical Limitations

We acknowledge several limitations to the quality of our data, as with any broad comparative analysis. Risk factor data was problematic or missing for certain viruses, e.g. natural transmission route for viruses only known to infect humans by accidental occupational exposure, and tissue tropism for viruses only known from serological evidence. However, the consistency of findings between alternative, stricter definitions of virulence and data subsets removing viruses with suspected data quality issues suggests scarcity of data does not bias our analyses.

Virulence also exhibits substantial variation at the sub-species level, i.e. between strains or variants. For example, severity of Lassa virus disease superficially varies with infection route and geography, though this appears to be driven by variation between genotypes [30]. Confirmatory analyses at a finer resolution would validate our identified risk factors, e.g. phylogenetic trait models of individual genera or species. Furthermore, clinical symptoms are also subject to traits of the host individual, e.g., immunocompetence, age, microbiome [31,32]. Our risk factor analysis brings a novel, top-down perspective on virulence at the broadest level, though caution must be exerted in extrapolating the risk factors we find to dynamics of specific infections.

### Implications for Public Health

The value of predictive modelling as an inexpensive and rapid tool for risk assessments during early emergence is increasingly recognised [16]. Instances where machine learning model predictions do not match outcomes could indicate likely candidates for outcome class changes, e.g. future reservoir hosts for zoonotic disease [33]. Severe virulence was predicted for one virus rated ‘nonsevere’ from literature protocols, *Usutu virus*, potentially suggesting the capability for more severe disease to be recognised in future.

However, our models have restricted function in predicting the virulence of a newly identified virus. Although taxonomy is easily accessible and applicable to give simple virulence estimates, the most informative non-taxonomic predictor, tissue tropism, is not likely to be known with confidence before clinical observations of virulence. One way to address this paucity of data lies in the potential predictability of tissue tropism from cell receptors, and more challengingly, cell receptors from viral sequence data [34], an increasingly accessible information source during early emergence following advances in genomic sequencing methods [35]. However, the exact links between tissue tropism, cell receptors, and sequences are currently a critical knowledge gap, but a potentially powerful focus for future predictive efforts. A further key area will be the possibility to directly infer virulence itself from other aspects of sequence data, e.g. genome composition biases, which have recently demonstrated the potential to predict reservoir host taxa and arthropod vectors via machine learning [36].

More widely, our analysis brings a novel focus that complements comparative models predicting other aspects of the emergence process, such as zoonotic transmission [8,9,27,33], propagation within humans [10,11] or geographic hotspots [37,38]. After continued calls for model-informed strategy, predictive studies are now beginning to shape surveillance and prevention with respect to emerging zoonoses [16,39], with virulence being been suggested as a factor to direct viral surveillance [40], albeit in non-human hosts. The virulence risk factors we identify suggest that broadly targeting direct contact or respiratory transmission interfaces within ecological systems and/or tailoring detection assays towards certain virus families (e.g. *Hantaviridae*) or tissues (e.g. neural tissue) could contribute to a viable strategy to detect future virulent zoonoses.

## Conclusion

This work adds to the comparative and predictive modelling efforts surrounding emerging infectious diseases. Here, we contribute a novel focus in ecological predictors of virulence of human RNA viruses, which can be combined in holistic frameworks with other models such as those predicting emergence dynamics. As a predictive model, the featured random forest offers valuable inference into the evolutionary determinants of virulence in newly emerging infections. We propose that future predictive studies and preparedness initiatives with respect to emerging diseases should carefully consider potential for human virulence.

## Materials and Methods

### Data Collection

For each of the 214 recognised human-infective RNA virus species following standardised data compilation efforts and critical assessment protocols [5], data on virulence and potential risk factors were collected via a systematic search and review of clinical and epidemiological literature. The following were consulted in turn: clinical virology textbooks [41–43]; references from the dataset described by [5]; literature searches using Google Scholar (search terms: 1) [virus name] AND human, 2) [virus name] AND human AND case, 3) [virus name] AND human AND [fatal* OR death], 4) [virus name] AND human AND [tropi* or isolat*]. Searches 3 and 4 were carried out only when fatality or tropism data respectively were not already found from previous sources. Data collection and virus name search terms included the full species name, any synonyms or subspecies (excluding vaccine strains) and the standard virus abbreviation as given by ICTV Online Virus Taxonomy [44].

Although many possible measurements of virulence have been proposed [45,46], even simple metrics like case fatality ratio (CFR) have not been calculated for the majority of human RNA virus species. Therefore, virulence was rated using a simple two-category measure of severity of typical disease in humans. We rated viruses as ‘severe’ if they firstly had ≥5% CFR where data was available (159/214 viruses including those with zero CFR), otherwise, we rated viruses as ‘severe’ if they had frequent reports of hospitalisation, were associated with significant morbidity from certain conditions (haemorrhagic fever, seizures/coma, cirrhosis, AIDS, hantavirus pulmonary syndrome, HTLV-associated myelopathy) or were explicitly described as “severe” or “causing severe disease” (S1 Table, S2 Table). We rated viruses as ‘nonsevere’ if none of these conditions were met. Note that this led to ‘nonsevere’ ratings for some viruses with clinically severe, but rare syndromes, e.g. Dengue virus can cause haemorrhagic dengue fever, though this is much rarer than typical acute dengue fever [41,42]. To address this, data were also collected on whether the virus has caused fatalities in vulnerable individuals (defined as age 16 and below or 60 and above, immunosuppressed, having co-morbidities, or otherwise cited as being ‘at-risk’ by sources for specific viruses) and in healthy adults, and whether any ‘nonsevere’ virus has atypically severe strains (for example, most infections with viruses within the species *Human enterovirus C* cause mild disease; however, poliovirus, which causes severe paralytic disease, is also classified under this species). These were examined both individually and within a composite six-rank system (S5 Table).

Data were compiled for four main risk factors: transmission route(s) and tissue tropisms, sourced from literature search exercises as described; and extent of human-to-human transmissibility and host range, sourced directly from [5]. Although evolutionary theories also predict virulence to vary with other traits, e.g. environmental survivability [47], paucity of data or nestedness within taxonomic family prevented their inclusion in our analysis. Transmission route was defined as the primary route the virus is transmitted by, classified as either vector-borne (excluding mechanical transmission), direct contact, faecal-oral or respiratory transmission. Tissue tropism was specified the primary organ system the virus typically infects or targets, classified as either neural, gastrointestinal, hepatic, respiratory, circulatory, vascular, or ‘systemic’ (primary tropism within multiple organ systems). We accepted isolation of the virus, viral proteins or genetic material, or diagnostic symptoms of the virus (such as characteristic histological damage) as evidence of infection within an organ system but did not accept generalised symptoms such as inflammation. However, many human viruses were isolated from blood with no further evidence of any specific tissue tropisms (n = 69). Therefore, we also included an additional ‘viraemia’ category in this variable to indicate only blood presence was known. Binary variables were also constructed denoting whether viruses were ever known to utilise a) more than one transmission route/tissue tropism, and b) each individual transmission route and tropism, including additional categories that were never among the primary routes/tropisms (food-borne and vertical transmission; renal, cardiac, joint, reproductive, sensory, skin, muscular and endocrine tropism).

Human-to-human transmissibility was specified using infectivity/transmissibility levels, based on previous conceptual models and a systematic compilation and review of evidence [4,5,12]. Level 2 denotes a virus capable of infecting humans but not transmitting between humans (R_0_ = 0), level 3 denotes a virus with limited human-to-human transmissibility (0 < R_0_ ≤ 1); and level 4 denotes a virus with sustained human-to-human transmissibility (R_0_ ≥ 1). Host range was specified as either ‘narrow’ (infection known only within humans or humans plus non-human primates) or ‘broad’ (infection known in mammals or animals beyond primates) [5]. Binary variables were also sourced as to whether infection was known within a) humans only, b) non-human primates, c) other mammals and d) birds. All virulence and risk factor data pertained to natural or unintentional artificially-acquired human infection only and data from intentional human infection, animal infection, and *in vitro* infection were not considered. Viral taxonomy was included in analyses by specifying both genome type and taxonomic family as predictors. All virulence and risk factor data are available via Figshare [48].

### Machine Learning Risk Factor Analysis

Firstly, the 212 retained virus species were split into a training set for model fitting and test set for model evaluation at an approximate 75:25 ratio using stratified random sampling based on taxonomic family and virulence rating. Fisher’s exact tests confirmed equal representation of families (p = 0.991) and virulence ratings (p > 0.999) between training and test data. Comparative risk factor analyses were firstly carried out by constructing a classification tree using the R package ‘rpart’ v4.1-11 [49]. Classification trees are a simple form of machine learning models that aim to optimally classify data points into their correct category of outcome variable based on a structure of binary predictor splits. Tree-based methods are well-suited for comparative analyses where confounding often results from taxonomic signal or suites of otherwise co-occurring traits as their high structure can intuitively fit complex non-linear interactions and local effects.

A tree model was fitted to the training set to predict virulence ratings by ‘recursive partitioning’, the repeated splitting of the dataset using every possible binary permutation of each predictor, and retaining the split that minimises the Gini impurity [50], defined as 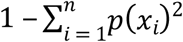 for outcome variable *x* with *n* possible ratings and *p*(*x*_*i*_) denoting proportion of data with rating *i*, which is equal to zero for perfectly separated data. To prevent overfitting, the tree was pruned back to the optimal branching size, taken as most common consensus size over 1000 repeats of 10-fold cross-validation. To validate the predictive power of the classification tree, predictions of virulence rating were generated when applied to the test set. Tree accuracy was then calculated comparing the proportion of correct predictions compared to literature-assigned ratings (assuming these to be 100% accurate as the ‘gold standard’ or ‘ground truth’). As virulence ratings were imbalanced (i.e. only a minority of viruses cause severe disease, so correct nonsevere classifications are likely to be achieved by chance), accuracy was directly compared to the null model, i.e. a model with no predictors that predicted ‘nonsevere’ for all viruses. Additional diagnostics of interest (sensitivity, specificity, negative predictive value, and True Skill Statistic [60]) were also obtained.

Although classification trees have the advantage of presenting an interpretable schematic of risk factor effects and directions, individual tree structures may be sensitive to particular data points and have no intuitive measures of uncertainty. Therefore, we constructed a random forest, an ensemble collection of a large number of bootstrapped classification trees [51]. Having many predictor variables compared to the relatively limited and fixed number of human-infective RNA virus species, random forests handle such ‘large p, small n’ data architecture much more easily than traditional regression frameworks [52]. Missing data in all predictors was imputed using the R package ‘missForest’ v1.4 [53]. Then, using the R package ‘randomForest’ v4.6-12 [53], a random forest was created containing 5000 individual trees, each built upon a bootstrapped sample of the training data and restricted to test a randomly selected subset of predictors (k = 5) at each split during construction and convergence confirmed by inspection. Predictive power of the random forest model was evaluated using the test set as for the classification tree and receiver operating characteristic curves were visualised and area under curves calculated to directly compare the two machine learning methodologies.

Due to their high structuring, random forest models cannot give a simple parametric predictor effect size and direction (e.g., an odds ratio). Instead, potential virulence risk factors were evaluated using two metrics: variable importance and partial dependence. Variable importance is calculated as the mean decrease in Gini impurity following tree splits on the predictor and can be considered as how informative the risk factor was towards correctly predicting virulence. Partial dependence is calculated as the mean relative change in log-odds of predicting severe virulence, which were converted to predicted probabilities of severity associated with each risk factor. Partial dependences describe marginal effects averaging across any influence of other predictors and as such, a single estimate may not reflect any complex risk factor interactions. Therefore, to test hypotheses regarding virulence risk factors, we present both random forest partial dependences and the less robust but more accessible single classification tree for its ease of interpretation in risk factor structure, and directly compare the statistical validity of both methods by plotting receiver operating characteristic curves. All modelling was carried out in R v 3.4.3 [54], with a supporting R script available via Figshare [48].

## Acknowledgements

We thank Jarrod Hadfield, Samantha Lycett, and Daniel Streicker for helpful discussion, and Alex Bhattacharya, Christopher McCaffery, David McCulloch, Conor O’Halloran, Claire Taylor and Feifei Zhang for assistance in data collection.

## Supporting Information Captions

**S1 Table. Virulence literature rating data for human RNA virus training dataset.**

Virulence data for the 181 virus species in the training set, ordered by genome type and taxonomy, including disease severity rating and supporting criteria for viruses rated ‘severe’, whether virus is known to have caused fatalities in vulnerable individuals and/or otherwise healthy adults, and whether virus is known to have ‘severe’ strains if species is rated ‘nonsevere’. CFR = Case fatality ratio, HPS = Hantavirus pulmonary syndrome, HFRS = Hantavirus haemorrhagic fever with renal syndrome, HTLV = Human T-lymphotropic virus, AIDS = Acquired immunodeficiency syndrome.

**S2 Table. Virulence literature rating data and predictions for human RNA virus test dataset.**

Virulence data for 31 virus species in the test set, ordered by genome type and taxonomy, whether virus is known to have caused fatalities in vulnerable individuals and/or otherwise healthy adults, and whether virus is known to have ‘severe’ strains if species is rated ‘nonsevere’. Both disease severity rating/supporting criteria following the literature protocol given in the main text, and predicted probability of severe disease from the random forest model are given. Bold type denotes where predictions do not match literature-based ratings. CFR = Case fatality ratio, HPS = Hantavirus pulmonary syndrome.

**S3 Table. Partial dependence from the random forest model for all predictor variables.**

Partial dependence given as marginal relative change in log-odds and predicted probability of classifying virulence as ‘severe’ from the random forest for all predictor variables.

**S4 Table. Diagnostics of random forest models using stringent data subsets.**

Predictive performance metrics of random forest models applied to datasets excluding viruses with low-certainty data (n denotes number of viruses excluded). In each case, data were randomly resampled using stratification upon taxonomic family and virulence rating, resulting in differing training and test sets from the main analysis. Otherwise, random forest methodology follows that of Materials & Methods.

**S5 Table. Six-rank system of classifying virulence for human RNA viruses.**

Six-rank system of classifying human RNA virus virulence with available data (specifically, severity rating from main text, fatalities in vulnerable individuals and healthy adults, and severe strains), along with example viruses and number of viruses fitting each exclusive rank’s criteria.

**S6 Table. Diagnostics of random forest models predicting alternative metrics of virulence.**

Predictive performance metrics of random forest models predicting alternative virulence measures using different two-category definitions of ‘severe’ (n denotes number of viruses considered ‘severe’ using that definition). Vulnerable individuals are defined as those age 16 and below, age 60 and above, immunosuppressed, having co-morbidities, or otherwise cited as being ‘at-risk’. Ranks follow those given in Table S5. Otherwise, random forest methodology follows that of Materials & Methods.

**S1 Fig. Variable importances from random forest models using stringent data subsets.**

Variable importance for virulence risk factors from random forest models applied to datasets excluding a) viruses only known to infect humans from serological evidence (n = 36), b) viruses with < 20 recognised human infections (n = 55), and c) viruses with poor data quality in at least one predictor (n = 71). Variable importance is calculated as the relative mean decrease in Gini impurity scaled against the most informative predictor within each model, alongside importances from the main analysis for comparison. ‘Tp’ denotes tissue tropism predictor, ‘Tr’ denotes transmission route predictor, ‘Tr level’ denotes level of human-to-human transmissibility, and ‘H’ denotes host range predictor.

**S2 Fig. Partial dependences from random forest models using stringent data subsets.**

Predicted probability of classifying virulence as ‘severe’ for each of the most informative risk factors from random forest models applied to datasets excluding a) viruses only known to infect humans from serological evidence (n = 36), b) viruses with < 20 recognised human infections (n = 55), and c) viruses with poor data quality in at least one predictor (n = 71), alongside predicted probabilities from the main analysis for comparison. Probabilities given are marginal, i.e. averaging over any effects of other predictors. As each data subset required random resampling of the training and test data, note that the raw prevalence of ‘severe’ virulence differed between each model (see S4 Table).

**S3 Fig. Variable importances from random forest models using stringent data subsets.**

Variable importance for virulence risk factors from random forest models predicting alternative virulence measures using different two-category definitions of ‘severe’, calculated as the relative mean decrease in Gini impurity scaled against the most informative predictor within each model, alongside importances from the main analysis for comparison. ‘Tp’ denotes tissue tropism predictor, ‘Tr’ denotes transmission route predictor, ‘Tr level’ denotes level of human-to-human transmissibility, and ‘H’ denotes host range predictor.

**S4 Fig. Partial dependences from random forest models using stringent data subsets.**

Predicted probability of classifying virulence as ‘severe’ in alternative virulence measures for each of the most informative risk factors from random forest models, alongside predicted probabilities from the main analysis for comparison. Probabilities given are marginal, i.e. averaging over any effects of other predictors. As each measurement used a different two-category definition of ‘severe’, note that the raw prevalence of ‘severe’ virulence differed between each model (see S6 Table).

## References

1. Morse SS. Factors in the emergence of infectious diseases. Emerg Infect Dis. 1995;1: 7–15.

2. Zaki AM, van Boheemen S, Bestebroer TM, Osterhaus ADME, Fouchier RAM. Isolation of a novel coronavirus from a man with pneumonia in Saudi Arabia. N Engl J Med. 2012;367: 1814–1820. doi:10.1056/NEJMoa1211721

3. Gatherer D, Kohl A. Zika virus: a previously slow pandemic spreads rapidly through the Americas. J Gen Virol. 2016;97: 269–73.

4. Woolhouse MEJ, Scott F, Hudson Z, Howey R, Chase-Topping M. Human viruses: discovery and emergence. Philos Trans R Soc B Biol Sci. 2012;367: 2864–2871. doi:10.1098/rstb.2011.0354

5. Woolhouse MEJ, Brierley L. Epidemiological characteristics of human-infective RNA viruses. Sci Data. 2018;5: 180017. doi:10.1038/sdata.2018.17

6. Woolhouse MEJ, Gowtage-Sequeria S. Host range and emerging and reemerging pathogens. Emerg Infect Dis. 2005;11: 1842–1847. doi:10.3201/eid1112.050997

7. Taylor LH, Latham SM, Woolhouse MEJ. Risk factors for human disease emergence. Philos Trans R Soc Lond B Biol Sci. 2001;356: 983–989.

8. Cleaveland S, Laurenson MK, Taylor LH. Diseases of humans and their domestic mammals: pathogen characteristics, host range and the risk of emergence. Philos Trans R Soc Lond B Biol Sci. 2001;356: 991–999.

9. Olival KJ, Hosseini PR, Zambrana-Torrelio C, Ross N, Bogich TL, Daszak P. Host and viral traits predict zoonotic spillover from mammals. Nature. 2017;546: 646–650. doi:10.1038/nature22975

10. Geoghegan JL, Senior AM, Giallonardo FD, Holmes EC. Virological factors that increase the transmissibility of emerging human viruses. Proc Natl Acad Sci. 2016;113: 4170–4175. doi:10.1073/pnas.1521582113

11. Johnson CK, Hitchens PL, Evans TS, Goldstein T, Thomas K, Clements A, et al. Spillover and pandemic properties of zoonotic viruses with high host plasticity. Sci Rep. 2015;5: 14830. doi:10.1038/srep14830

12. Woolhouse MEJ, Brierley L, McCaffery C, Lycett S. Assessing the Epidemic Potential of RNA and DNA Viruses. Emerg Infect Dis. 2016;22: 2037–2044. doi:10.3201/eid2212.160123

13. Feldmann H, Geisbert TW. Ebola haemorrhagic fever. The Lancet. 2011;377: 849–862. doi:10.1016/S0140-6736(10)60667-8

14. Focosi D, Maggi F. Estimates of Ebola virus case-fatality ratio in the 2014 West African outbreak. Clin Infect Dis. 2015;60: 829. doi:10.1093/cid/ciu921

15. Morikawa S, Saijo M, Kurane I. Current knowledge on lower virulence of Reston Ebola virus. Comp Immunol Microbiol Infect Dis. 2007;30: 391–398. doi:10.1016/j.cimid.2007.05.005

16. Morse SS, Mazet JA, Woolhouse MEJ, Parrish CR, Carroll D, Karesh WB, et al. Prediction and prevention of the next pandemic zoonosis. The Lancet. 2012;380: 1956–1965. doi:10.1016/S0140-6736(12)61684-5

17. Walther BA, Ewald PW. Pathogen survival in the external environment and the evolution of virulence. Biol Rev. 2004;79: 849–869. doi:10.1017/S1464793104006475

18. Ewald PW. Host-parasite relations, vectors, and the evolution of disease severity. Annu Rev Ecol Syst. 1983;14: 465–485. doi:10.2307/2096982

19. Leggett HC, Buckling A, Long GH, Boots M. Generalism and the evolution of parasite virulence. Trends Ecol Evol. 2013;28: 592–596. doi:10.1016/j.tree.2013.07.002

20. Anderson RM, May RM. Coevolution of hosts and parasites. Parasitology. 1982;85: 411–426. doi:10.1017/S0031182000055360

21. Bremermann HJ, Pickering J. A game-theoretical model of parasite virulence. J Theor Biol. 1983;100: 411–426. doi:10.1016/0022-5193(83)90438-1

22. Alizon S, Hurford A, Mideo N, Van Baalen M. Virulence evolution and the trade-off hypothesis: history, current state of affairs and the future. J Evol Biol. 2009;22: 245–259. doi:10.1111/j.1420-9101.2008.01658.x

23. Ebert D, Bull JJ. Challenging the trade-off model for the evolution of virulence: is virulence management feasible? Trends Microbiol. 2003;11: 15–20. doi:10.1016/S0966-842X(02)00003-3

24. Bull JJ. Perspective: virulence. Evolution. 1994;48: 1423–1437. doi:10.2307/2410237

25. Levin B., Svanborg Edén C. Selection and evolution of virulence in bacteria: an ecumenical excursion and modest suggestion. Parasitology. 1990;100: S103–S115. doi:10.1017/S0031182000073054

26. Longdon B, Hadfield JD, Day JP, Smith SCL, McGonigle JE, Cogni R, et al. The causes and consequences of changes in virulence following pathogen host shifts. PLoS Pathog. 2015;11: e1004728. doi:10.1371/journal.ppat.1004728

27. Pedersen AB, Davies TJ. Cross-species pathogen transmission and disease emergence in primates. EcoHealth. 2009;6: 496–508.

28. Levin BR, Bull JJ. Short-sighted evolution and the virulence of pathogenic microorganisms. Trends Microbiol. 1994;2: 76–81. doi:10.1016/0966-842X(94)90538-X

29. Taber SW, Pease CM. Paramyxovirus phylogeny: tissue tropism evolves slower than host specificity. Evolution. 1990;44: 435–438. doi:10.2307/2409419

30. Howard CR. Arenaviruses. In: Zuckerman AJ, Banatvala JE, Schoub BD, Griffiths PD, Mortimer P, editors. Principles and practice of clinical virology. John Wiley & Sons, Ltd; 2009. pp. 733–754.

31. Mackinnon MJ, Gandon S, Read AF. Virulence evolution in response to vaccination: The case of malaria. Vaccine. 2008;26, Supplement 3: C42–C52. doi:10.1016/j.vaccine.2008.04.012

32. Franco DJ, Vago AR, Chiari E, Meira FCA, Galvão LMC, Machado CRS. Trypanosoma cruzi: mixture of two populations can modify virulence and tissue tropism in rat. Exp Parasitol. 2003;104: 54–61.

33. Han BA, Schmidt JP, Bowden SE, Drake JM. Rodent reservoirs of future zoonotic diseases. Proc Natl Acad Sci. 2015;112: 7039–7044. doi:10.1073/pnas.1501598112

34. Woolhouse M. Sources of human viruses. Science. 2018;362: 524–525. doi:10.1126/science.aav4265

35. Woolhouse MEJ, Rambaut A, Kellam P. Lessons from Ebola: Improving infectious disease surveillance to inform outbreak management. Sci Transl Med. 2015;7: 307rv5. doi:10.1126/scitranslmed.aab0191

36. Babayan SA, Orton RJ, Streicker DG. Predicting reservoir hosts and arthropod vectors from evolutionary signatures in RNA virus genomes. Science. 2018;362: 577–580. doi:10.1126/science.aap9072

37. Jones KE, Patel NG, Levy MA, Storeygard A, Balk D, Gittleman JL, et al. Global trends in emerging infectious diseases. Nature. 2008;451: 990–993.

38. Allen T, Murray KA, Zambrana-Torrelio C, Morse SS, Rondinini C, Marco MD, et al. Global hotspots and correlates of emerging zoonotic diseases. Nat Commun. 2017;8: 1124. doi:10.1038/s41467-017-00923-8

39. Daszak P. A call for “smart surveillance”: a lesson learned from H1N1. EcoHealth. 2009;6: 1–2.

40. Levinson J, Bogich TL, Olival KJ, Epstein JH, Johnson CK, Karesh W, et al. Targeting surveillance for zoonotic virus discovery. Emerg Infect Dis. 2013;19: 743–747. doi:10.3201/eid1905.121042

41. Knipe DM, Howley PM. Fields virology, 5th Edition. Lippincott Williams & Wilkins; 2007.

42. Zuckerman AJ, Banatvala JE, Griffiths P, Schoub B, Mortimer P. Principles and practice of clinical virology. John Wiley & Sons; 2009.

43. Richman DD, Whitley RJ, Hayden FG. Clinical virology. John Wiley & Sons; 2009.

44. ICTV. The Classification and Nomenclature of Viruses. The Online (10th) Report of the ICTV. [Internet]. 2017. Available: https://talk.ictvonline.org/ictv-reports/ictv_online_report/

45. Nathanson N, Gonzalez-Scarano F, Nathanson N. Viral virulence. Viral Pathogenesis and Immunity. Academic Press; 2007. pp. 113–129.

46. Day T. On the evolution of virulence and the relationship between various measures of mortality. Proc R Soc B Biol Sci. 2002;269: 1317–1323. doi:10.1098/rspb.2002.2021

47. Bonhoeffer S, Lenski RE, Ebert D. The curse of the pharaoh: the evolution of virulence in pathogens with long living propagules. Proc R Soc Lond B Biol Sci. 1996;263: 715–721.

48. Brierley L, Pedersen A, Woolhouse M. Data and supporting R script for: Tissue Tropism and Transmission Ecology Predict Virulence of Human RNA Viruses [Internet]. figshare. 2019. doi:10.6084/m9.figshare.7406441.v1

49. Therneau TM, Atkinson B, Ripley B. rpart: Recursive partitioning and regression Trees. R package version 4. 1–8. 2014;

50. De’ath G, Fabricius KE. Classification and regression trees: a powerful yet simple technique for ecological data analysis. Ecology. 2000;81: 3178–3192.

51. Breiman L. Random forests. Mach Learn. 2001;45: 5–32. doi:10.1023/A:1010933404324

52. Genuer R, Poggi J-M, Tuleau C. Random Forests: some methodological insights. ArXiv Prepr ArXiv08113619. 2008;

53. Stekhoven DJ, Bühlmann P. MissForest—non-parametric missing value imputation for mixed-type data. Bioinformatics. 2012;28: 112–118. doi:10.1093/bioinformatics/btr597

54. R Development Core Team. R: A language and environment for statistical computing. R Foundation for Statistical Computing, Vienna, Austria. http://www.R-project.org; 2011.

