## Supplementary figures and images for "Tissue Tropism and Transmission Ecology Predict Virulence of Human RNA Viruses"

### Supplemental Figure S1

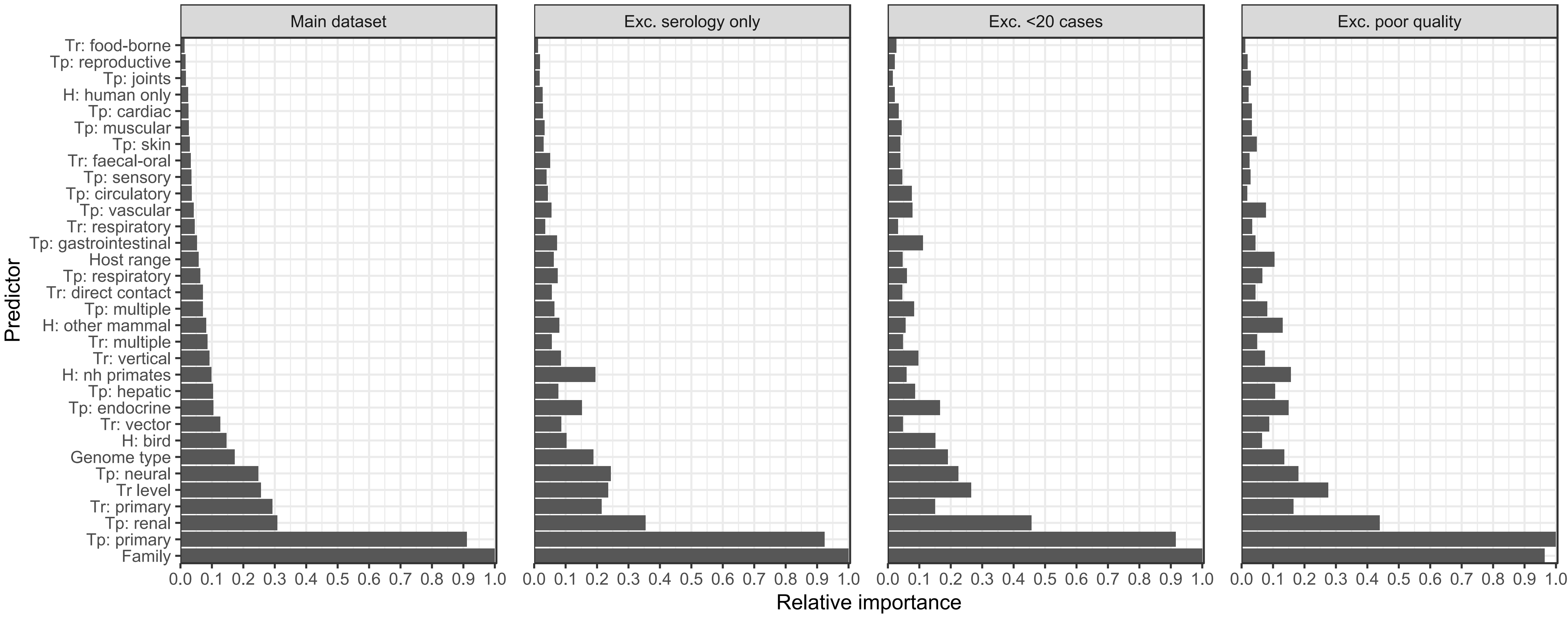

### Supplemental Figure S2

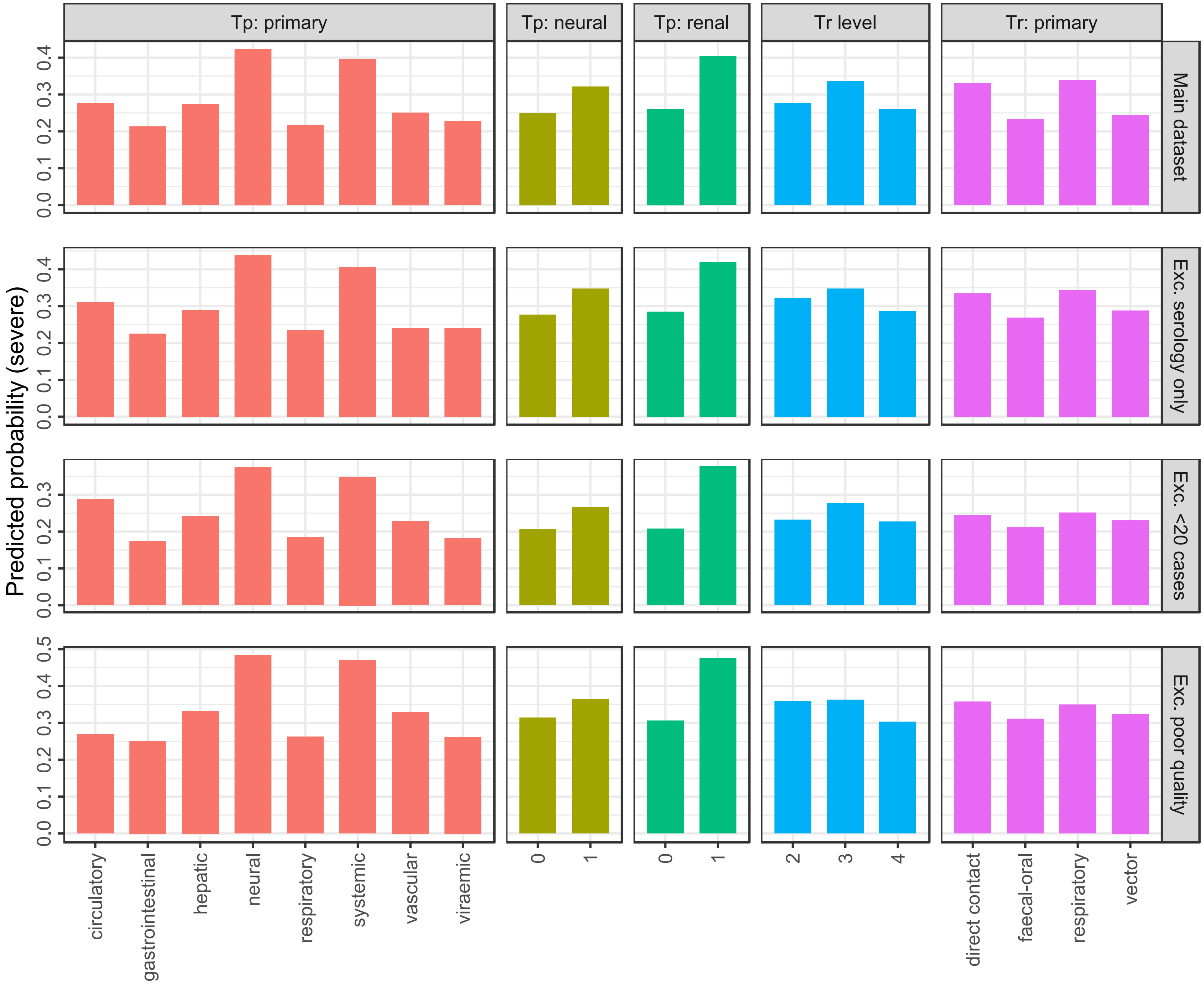

### Supplemental Figure S3

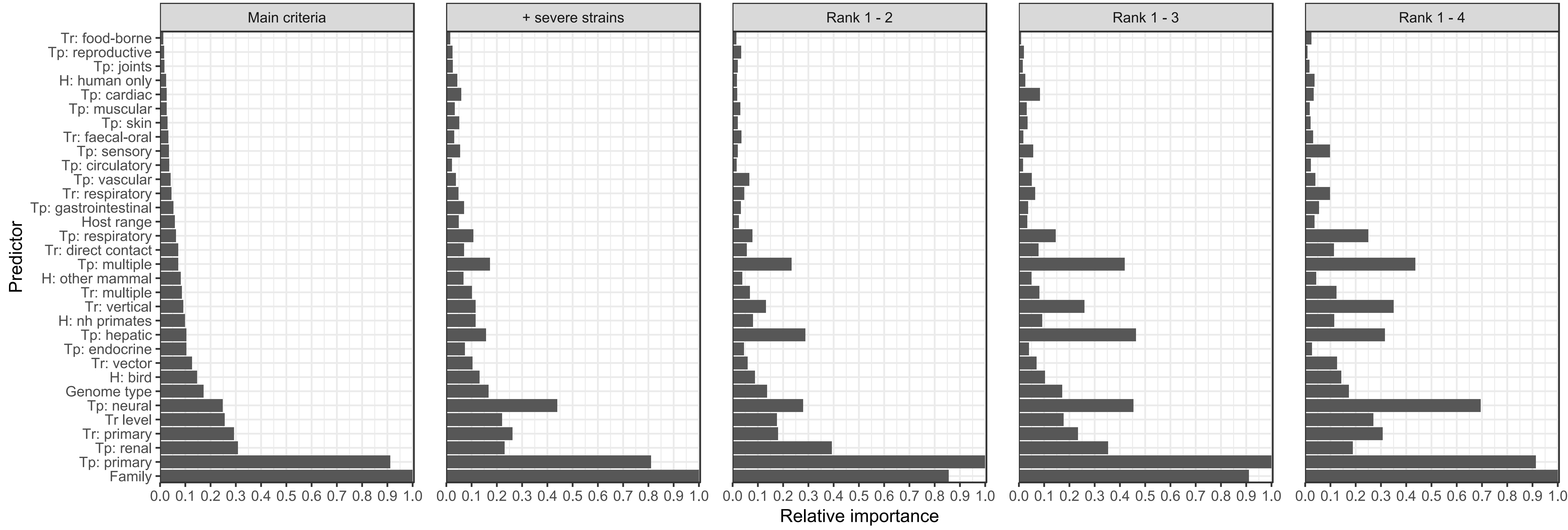

### Supplemental Figure S4

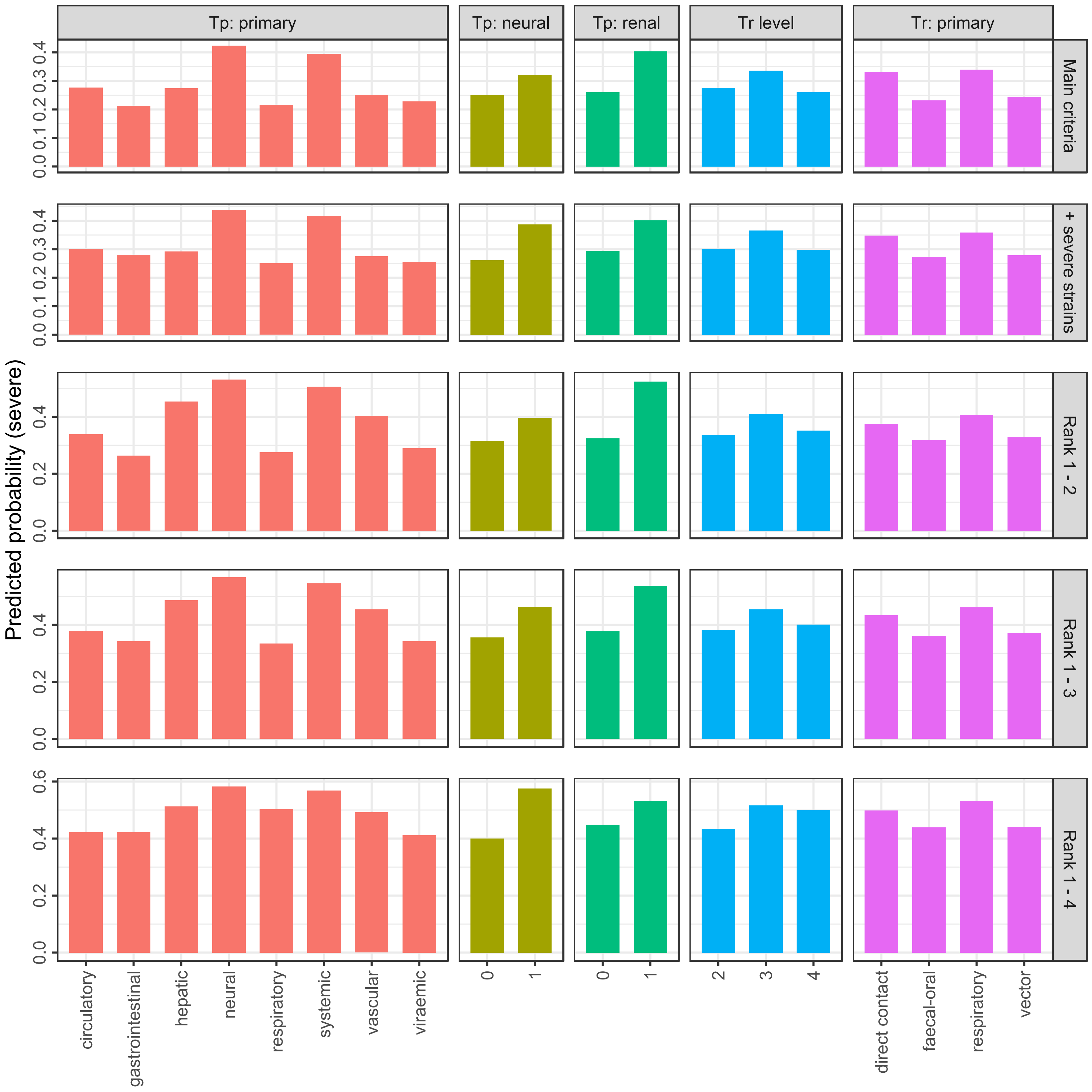
