## Supplemental Table S1 for "Tissue Tropism and Transmission Ecology Predict Virulence of Human RNA Viruses"

| [1]Family | Genus | Species | Severity rating (literature protocol) | Severity notes | Fatalities (vulnerable) | Fatalities (healthy adults) | Severe strains | References |
| --- | --- | --- | --- | --- | --- | --- | --- | --- |
| *-ssRNA viruses* | | |  |  |  |  |  |  |
| *Arenaviridae* | *Mammarenavirus* | *Chapare mammarenavirus* | severe | Haemorrhagic fever, CFR 100% (1 case) | 1 | 1 | N/A | [1–3] |
| *Arenaviridae* | *Mammarenavirus* | *Guanarito mammarenavirus* | severe | Haemorrhagic fever, CFR 25% | 1 | 1 | N/A | [1,4] |
| *Arenaviridae* | *Mammarenavirus* | *Junín mammarenavirus* | severe | Haemorrhagic fever, CFR 15-30% | 1 | 1 | N/A | [1,4] |
| *Arenaviridae* | *Mammarenavirus* | *Lujo mammarenavirus* | severe | Haemorrhagic fever, CFR 80% (5 cases) | 1 | 1 | N/A | [5,6] |
| *Arenaviridae* | *Mammarenavirus* | *Lymphocytic choriomeningitis mammarenavirus* | nonsevere |  | 1 | 1 | 0 | [1,2,4] |
| *Arenaviridae* | *Mammarenavirus* | *Machupo mammarenavirus* | severe | Haemorrhagic fever, CFR 5-35% | 1 | 1 | N/A | [1,2,4] |
| *Arenaviridae* | *Mammarenavirus* | *Mobala mammarenavirus* | nonsevere |  | 0 | 0 | 0 | [7] |
| *Arenaviridae* | *Mammarenavirus* | *Pichindé mammarenavirus* | nonsevere |  | 0 | 0 | 0 | [4] |
| *Arenaviridae* | *Mammarenavirus* | *Sabiá mammarenavirus* | severe | Haemorrhagic fever, CFR 33% (3 cases) | 1 | 1 | N/A | [1,2,4] |
| *Bornaviridae* | *Bornavirus* | *Mammalian 1 bornavirus* | nonsevere |  | 0 | 0 | 0 | [4] |
| *Bornaviridae* | *Bornavirus* | *Mammalian 2 bornavirus* | severe | CFR 100% (3 cases) | 1 | 0 | N/A | [8] |
| *Filoviridae* | *Ebolavirus* | *Bundibugyo ebolavirus* | severe | Haemorrhagic fever, CFR 35% | 1 | 1 | N/A | [9,10] |
| *Filoviridae* | *Ebolavirus* | *Reston ebolavirus* | nonsevere |  | 0 | 0 | 0 | [2,4] |
| *Filoviridae* | *Ebolavirus* | *Sudan ebolavirus* | severe | Haemorrhagic fever, CFR 41-65% | 1 | 1 | N/A | [1,4] |
| *Filoviridae* | *Ebolavirus* | *Tai forest ebolavirus* | severe | Haemorrhagic fever | 0 | 0 | N/A | [1,4] |
| *Filoviridae* | *Marburgvirus* | *Marburg marburgvirus* | severe | Haemorrhagic fever, CFR 23-90% | 1 | 1 | N/A | [4] |
| *Hantaviridae* | *Orthohantavirus* | *Andes orthohantavirus* | severe | HPS | 1 | 1 | N/A | [2] |
| *Hantaviridae* | *Orthohantavirus* | *Bayou orthohantavirus* | severe | HPS | 1 | 0 | N/A | [2,11] |
| *Hantaviridae* | *Orthohantavirus* | *Black creek canal orthohantavirus* | severe | HPS | 0 | 0 | N/A | [2] |
| *Hantaviridae* | *Orthohantavirus* | *Dobrava-Belgrade orthohantavirus* | severe | HFRS, CFR 5-35% | 1 | 1 | N/A | [2,4] |
| *Hantaviridae* | *Orthohantavirus* | *Hantaan orthohantavirus* | severe | HFRS, CFR 5-15% | 1 | 1 | N/A | [4] |
| *Hantaviridae* | *Orthohantavirus* | *Sangassou orthohantavirus* | severe | HFRS | 0 | 0 | N/A | [12] |
| *Hantaviridae* | *Orthohantavirus* | *Seoul orthohantavirus* | nonsevere |  | 1 | 1 | 0 | [4] |
| *Hantaviridae* | *Orthohantavirus* | *Sin Nombre orthohantavirus* | severe | HPS, CFR 32-75% | 1 | 1 | N/A | [1,4] |
| *Hantaviridae* | *Orthohantavirus* | *Thailand orthohantavirus* | nonsevere |  | 0 | 0 | 0 | [13] |
| *Hantaviridae* | *Orthohantavirus* | *Thottapalayam orthohantavirus* | severe | Only known case required hospitalisation | 0 | 0 | N/A | [14] |
| *Hantaviridae* | *Orthohantavirus* | *Tula orthohantavirus* | nonsevere |  | 0 | 0 | 0 | [15] |
| *Nairoviridae* | *Orthonairovirus* | *Crimean-Congo haemorrhagic fever orthonairovirus* | severe | Haemorrhagic fever, CFR 30% | 1 | 1 | N/A | [1,4] |
| *Nairoviridae* | *Orthonairovirus* | *Dugbe orthonairovirus* | nonsevere |  | 0 | 0 | 0 | [2] |
| *Nairoviridae* | *Orthonairovirus* | *Thiafora orthonairovirus* | nonsevere |  | 0 | 0 | 0 | [16,17] |
| *Orthomyxoviridae* | *Influenzavirus A* | *Influenza A virus* | nonsevere |  | 1 | 1 | 1 | [1,4] |
| *Orthomyxoviridae* | *Influenzavirus B* | *Influenza B virus* | nonsevere |  | 1 | 1 | 0 | [4] |
| *Orthomyxoviridae* | *Influenzavirus C* | *Influenza C virus* | nonsevere |  | 0 | 0 | 0 | [4] |
| *Orthomyxoviridae* | *Thogotovirus* | *Dhori virus* | nonsevere |  | 0 | 0 | 0 | [18] |
| *Orthomyxoviridae* | *Thogotovirus* | *Thogoto virus* | nonsevere |  | 1 | 0 | 0 | [19] |
| *Paramyxoviridae* | *Avulavirus* | *Avian avulavirus 1* | nonsevere |  | 1 | 0 | 0 | [1,20] |
| *Paramyxoviridae* | *Henipavirus* | *Hendra henipavirus* | severe | CFR 57% (7 cases) | 1 | 1 | N/A | [4,21] |
| *Paramyxoviridae* | *Henipavirus* | *Nipah henipavirus* | severe | CFR 10-75% | 1 | 1 | N/A | [1,4] |
| *Paramyxoviridae* | *Morbillivirus* | *Canine morbillivirus* | nonsevere |  | 0 | 0 | 0 | [22] |
| *Paramyxoviridae* | *Morbillivirus* | *Measles morbillivirus* | nonsevere |  | 1 | 1 | 0 | [2,4] |
| *Paramyxoviridae* | *Respirovirus* | *Human respirovirus 1* | nonsevere |  | 0 | 0 | 0 | [4] |
| *Paramyxoviridae* | *Respirovirus* | *Human respirovirus 3* | nonsevere |  | 1 | 0 | 0 | [4] |
| *Paramyxoviridae* | *Rubulavirus* | *Human rubulavirus 2* | nonsevere |  | 0 | 0 | 0 | [4] |
| *Paramyxoviridae* | *Rubulavirus* | *Human rubulavirus 4* | nonsevere |  | 0 | 0 | 0 | [4] |
| *Paramyxoviridae* | *Rubulavirus* | *Mammalian rubulavirus 5* | nonsevere |  | 0 | 0 | 0 | [4,23] |
| *Paramyxoviridae* | *Rubulavirus* | *Mumps rubulavirus* | nonsevere |  | 1 | 1 | 0 | [4] |
| *Paramyxoviridae* | *Rubulavirus* | *Sosuga rubulavirus* | severe | Single known case required hospital admission | 0 | 0 | N/A | [24] |
| *Paramyxoviridae* | *Rubulavirus* | *Tioman rubulavirus* | nonsevere |  | 0 | 0 | 0 | [25] |
| *Peribunyaviridae* | *Orthobunyavirus* | *Bunyamwera orthobunyavirus* | nonsevere |  | 0 | 0 | 0 | [2,4] |
| *Peribunyaviridae* | *Orthobunyavirus* | *Bwamba orthobunyavirus* | nonsevere |  | 0 | 0 | 0 | [2] |
| *Peribunyaviridae* | *Orthobunyavirus* | *California encephalitis orthobunyavirus* | severe | High frequency of severe symptoms (seizures, coma) | 1 | 1 | N/A | [4] |
| *Peribunyaviridae* | *Orthobunyavirus* | *Caraparu orthobunyavirus* | nonsevere |  | 0 | 0 | 0 | [2] |
| *Peribunyaviridae* | *Orthobunyavirus* | *Catu orthobunyavirus* | nonsevere |  | 0 | 0 | 0 | [2] |
| *Peribunyaviridae* | *Orthobunyavirus* | *Guama orthobunyavirus* | nonsevere |  | 0 | 0 | 0 | [2] |
| *Peribunyaviridae* | *Orthobunyavirus* | *Madrid orthobunyavirus* | nonsevere |  | 0 | 0 | 0 | [2,4,26] |
| *Peribunyaviridae* | *Orthobunyavirus* | *Nyando orthobunyavirus* | nonsevere |  | 0 | 0 | 0 | [2,27] |
| *Peribunyaviridae* | *Orthobunyavirus* | *Oropouche orthobunyavirus* | nonsevere |  | 0 | 0 | 0 | [1,4] |
| *Peribunyaviridae* | *Orthobunyavirus* | *Patois orthobunyavirus* | nonsevere |  | 0 | 0 | 0 | [28] |
| *Peribunyaviridae* | *Orthobunyavirus* | *Shuni orthobunyavirus* | nonsevere |  | 0 | 0 | 0 | [2] |
| *Peribunyaviridae* | *Orthobunyavirus* | *Tacaiuma orthobunyavirus* | nonsevere |  | 0 | 0 | 0 | [2,29] |
| *Peribunyaviridae* | *Orthobunyavirus* | *Wyeomyia orthobunyavirus* | nonsevere |  | 0 | 0 | 0 | [2,30] |
| *Phenuiviridae* | *Phlebovirus* | *Candiru phlebovirus* | nonsevere |  | 0 | 0 | 0 | [2,31] |
| *Phenuiviridae* | *Phlebovirus* | *Punta Toro phlebovirus* | nonsevere |  | 0 | 0 | 0 | [1,2] |
| *Phenuiviridae* | *Phlebovirus* | *Rift Valley fever phlebovirus* | nonsevere |  | 1 | 1 | 0 | [2] |
| *Phenuiviridae* | *Phlebovirus* | *Sandfly fever Naples phlebovirus* | nonsevere |  | 0 | 0 | 0 | [1,2] |
| *Phenuiviridae* | *Phlebovirus* | *SFTS phlebovirus* | severe | CFR 15% | 1 | 1 | N/A | [32–35] |
| *Pneumoviridae* | *Metapneumovirus* | *Avian metapneumovirus* | nonsevere |  | 0 | 0 | 0 | [36] |
| *Pneumoviridae* | *Metapneumovirus* | *Human metapneumovirus* | nonsevere |  | 1 | 0 | 0 | [4] |
| *Pneumoviridae* | *Orthopneumovirus* | *Human orthopneumovirus* | nonsevere |  | 1 | 0 | 0 | [2,4] |
| *Rhabdoviridae* | *Lyssavirus* | *Australian bat lyssavirus* | severe | Rabies-like disease, CFR 100% (3 cases) | 1 | 1 | N/A | [2] |
| *Rhabdoviridae* | *Lyssavirus* | *Duvenhage lyssavirus* | severe | Rabies-like disease, CFR 75% (4 cases) | 1 | 1 | N/A | [1,2] |
| *Rhabdoviridae* | *Lyssavirus* | *European bat 1 lyssavirus* | severe | Rabies-like disease, CFR 100% (3 cases) | 1 | 0 | N/A | [2] |
| *Rhabdoviridae* | *Lyssavirus* | *European bat 2 lyssavirus* | severe | Rabies-like disease, CFR 20-50% | 1 | 1 | N/A | [1,2] |
| *Rhabdoviridae* | *Lyssavirus* | *Mokola lyssavirus* | severe | Neurologic disease different to classical rabies, CFR 33% (3 cases) | 1 | 0 | N/A | [2,37] |
| *Rhabdoviridae* | *Lyssavirus* | *Rabies lyssavirus* | severe | Rabies disease, CFR 60% | 1 | 1 | N/A | [4] |
| *Rhabdoviridae* | *Tibrovirus* | *Bas-Congo tibrovirus* | severe | Haemorrhagic fever, CFR 67% (3 cases) | 1 | 0 | N/A | [38] |
| *Rhabdoviridae* | *Tibrovirus* | *Ekpoma 1 tibrovirus* | nonsevere |  | 0 | 0 | 0 | [39] |
| *Rhabdoviridae* | *Tibrovirus* | *Ekpoma 2 tibrovirus* | nonsevere |  | 0 | 0 | 0 | [39] |
| *Rhabdoviridae* | *Vesiculovirus* | *Alagoas vesiculovirus* | nonsevere |  | 0 | 0 | 0 | [1,2,40] |
| *Rhabdoviridae* | *Vesiculovirus* | *Cocal vesiculovirus* | nonsevere |  | 0 | 0 | 0 | [41] |
| *Rhabdoviridae* | *Vesiculovirus* | *Indiana vesiculovirus* | nonsevere |  | 0 | 0 | 0 | [1,2,40] |
| *Rhabdoviridae* | *Vesiculovirus* | *Maraba vesiculovirus* | nonsevere |  | 0 | 0 | 0 | [4,42] |
| *Rhabdoviridae* | *Vesiculovirus* | *New Jersey vesiculovirus* | nonsevere |  | 0 | 0 | 0 | [1,2,40] |
| *Rhabdoviridae* | *Vesiculovirus* | *Piry vesiculovirus* | nonsevere |  | 0 | 0 | 0 | [1,43] |
| *+ssRNA viruses* | |  |  |  |  |  |  |  |
| *Astroviridae* | *Mamastrovirus* | *Mamastrovirus 1* | nonsevere |  | 1 | 0 | 0 | [2,4,44] |
| *Astroviridae* | *Mamastrovirus* | *Mamastrovirus 8* | nonsevere |  | 1 | 0 | 1 | [45–48] |
| *Astroviridae* | *Mamastrovirus* | *Mamastrovirus 9* | nonsevere |  | 0 | 0 | 0 | [45,49] |
| *Caliciviridae* | *Norovirus* | *Norwalk virus* | nonsevere |  | 1 | 0 | 0 | [1,4] |
| *Caliciviridae* | *Sapovirus* | *Sapporo virus* | nonsevere |  | 0 | 0 | 0 | [4] |
| *Caliciviridae* | *Vesivirus* | *Vesicular exanthema of swine virus* | nonsevere |  | 0 | 0 | 0 | [50] |
| *Coronaviridae* | *Alphacoronavirus* | *Alphacoronavirus 1* | nonsevere |  | 0 | 0 | 0 | [51] |
| *Coronaviridae* | *Alphacoronavirus* | *Human coronavirus 229E* | nonsevere |  | 1 | 0 | 0 | [2,4,52] |
| *Coronaviridae* | *Alphacoronavirus* | *Human coronavirus NL63* | nonsevere |  | 1 | 0 | 0 | [1,2,53] |
| *Coronaviridae* | *Betacoronavirus* | *Betacoronavirus 1* | nonsevere |  | 1 | 0 | 0 | [2,4,54] |
| *Coronaviridae* | *Betacoronavirus* | *Human coronavirus HKU1* | nonsevere |  | 0 | 0 | 0 | [2,4] |
| *Coronaviridae* | *Betacoronavirus* | *Middle East respiratory syndrome-related coronavirus* | severe | CFR 27-56% | 1 | 1 | N/A | [55–57] |
| *Coronaviridae* | *Betacoronavirus* | *Severe acute respiratory syndrome-related coronavirus* | severe | CFR 9-12% | 1 | 1 | N/A | [2,4] |
| *Coronaviridae* | *Torovirus* | *Human torovirus* | nonsevere |  | 0 | 0 | 0 | [1,2] |
| *Flaviviridae* | *Flavivirus* | *Aroa virus* | nonsevere |  | 0 | 0 | 0 | [58] |
| *Flaviviridae* | *Flavivirus* | *Bagaza virus* | nonsevere |  | 0 | 0 | 0 | [59] |
| *Flaviviridae* | *Flavivirus* | *Banzi virus* | nonsevere |  | 0 | 0 | 0 | [1,2,4] |
| *Flaviviridae* | *Flavivirus* | *Cacipacore virus* | severe | Only known case required hospitalisation | 0 | 0 | N/A | [60] |
| *Flaviviridae* | *Flavivirus* | *Dengue virus* | nonsevere |  | 1 | 1 | 0 | [2,4] |
| *Flaviviridae* | *Flavivirus* | *Edge Hill virus* | nonsevere |  | 0 | 0 | 0 | [1,61] |
| *Flaviviridae* | *Flavivirus* | *Gadgets Gully virus* | nonsevere |  | 0 | 0 | 0 | [4] |
| *Flaviviridae* | *Flavivirus* | *Ilheus virus* | severe | Described as potentially severe, one subspecies (Rocio virus) has CFR 10% | 1 | 1 | N/A | [1] |
| *Flaviviridae* | *Flavivirus* | *Japanese encephalitis virus* | severe | CFR 5-40% | 1 | 1 | N/A | [4] |
| *Flaviviridae* | *Flavivirus* | *Kokobera virus* | nonsevere |  | 0 | 0 | 0 | [1,4] |
| *Flaviviridae* | *Flavivirus* | *Kyasanur forest disease virus* | severe | CFR 3-25% | 1 | 1 | N/A | [1,2,4] |
| *Flaviviridae* | *Flavivirus* | *Louping ill virus* | nonsevere |  | 0 | 0 | 0 | [1,2] |
| *Flaviviridae* | *Flavivirus* | *Murray Valley encephalitis virus* | severe | CFR 18-42%, frequent neurologic sequelae | 1 | 1 | N/A | [1,2,4] |
| *Flaviviridae* | *Flavivirus* | *Omsk hemorrhagic fever virus* | nonsevere |  | 1 | 1 | 0 | [62] |
| *Flaviviridae* | *Flavivirus* | *Powassan virus* | severe | CFR 10-15%, frequent neurologic sequelae | 1 | 0 | N/A | [1,2,4] |
| *Flaviviridae* | *Flavivirus* | *St. Louis encephalitis virus* | severe | CFR 7-30% | 1 | 1 | N/A | [4] |
| *Flaviviridae* | *Flavivirus* | *Tick-borne encephalitis virus* | severe | CFR 1-54% | 1 | 1 | N/A | [2,4] |
| *Flaviviridae* | *Flavivirus* | *Uganda S virus* | nonsevere |  | 0 | 0 | 0 | [2,63] |
| *Flaviviridae* | *Flavivirus* | *Wesselsbron virus* | nonsevere |  | 0 | 0 | 0 | [1,4] |
| *Flaviviridae* | *Flavivirus* | *West Nile virus* | nonsevere |  | 1 | 1 | 0 | [4] |
| *Flaviviridae* | *Flavivirus* | *Zika virus* | nonsevere |  | 0 | 0 | 0 | [2,4] |
| *Flaviviridae* | *Hepacivirus* | *Hepacivirus C* | severe | Chronic disease including cirrhosis | 1 | 1 | N/A | [4] |
| *Flaviviridae* | *Pegivirus* | *Pegivirus A* | nonsevere |  | 0 | 0 | 0 | [64] |
| *Flaviviridae* | *Pestivirus* | *Bovine viral diarrhea virus 1* | nonsevere |  | 0 | 0 | 0 | [65] |
| *Hepeviridae* | *Orthohepevirus* | *Orthohepevirus A* | nonsevere |  | 1 | 1 | 0 | [1,4] |
| *Picornaviridae* | *Aphthovirus* | *Equine rhinitis A virus* | nonsevere |  | 0 | 0 | 0 | [66] |
| *Picornaviridae* | *Aphthovirus* | *Foot-and-mouth disease virus* | nonsevere |  | 0 | 0 | 0 | [67] |
| *Picornaviridae* | *Cardiovirus* | *Cardiovirus A* | nonsevere |  | 0 | 0 | 0 | [68] |
| *Picornaviridae* | *Cardiovirus* | *Cardiovirus B* | nonsevere |  | 1 | 1 | 1 | [4,69–71] |
| *Picornaviridae* | *Cosavirus* | *Cosavirus A* | nonsevere |  | 0 | 0 | 0 | [72–75] |
| *Picornaviridae* | *Cosavirus* | *Cosavirus B* | nonsevere |  | 0 | 0 | 0 | [72–74] |
| *Picornaviridae* | *Cosavirus* | *Cosavirus E* | nonsevere |  | 0 | 0 | 0 | [73,76,77] |
| *Picornaviridae* | *Cosavirus* | *Cosavirus F* | nonsevere |  | 0 | 0 | 0 | [73] |
| *Picornaviridae* | *Enterovirus* | *Enterovirus A* | nonsevere |  | 1 | 1 | 1 | [1,78] |
| *Picornaviridae* | *Enterovirus* | *Enterovirus B* | nonsevere |  | 1 | 1 | 1 | [1,4] |
| *Picornaviridae* | *Enterovirus* | *Enterovirus C* | nonsevere |  | 1 | 1 | 1 | [1,4] |
| *Picornaviridae* | *Enterovirus* | *Rhinovirus A* | nonsevere |  | 1 | 0 | 0 | [1,4,79] |
| *Picornaviridae* | *Enterovirus* | *Rhinovirus B* | nonsevere |  | 0 | 0 | 0 | [1,4] |
| *Picornaviridae* | *Enterovirus* | *Rhinovirus C* | nonsevere |  | 1 | 0 | 0 | [1,4,79] |
| *Picornaviridae* | *Erbovirus* | *Erbovirus A* | nonsevere |  | 0 | 0 | 0 | [80] |
| *Picornaviridae* | *Hepatovirus* | *Hepatovirus A* | nonsevere |  | 1 | 1 | 0 | [1,4] |
| *Picornaviridae* | *Kobuvirus* | *Aichivirus A* | nonsevere |  | 0 | 0 | 0 | [81,82] |
| *Picornaviridae* | *Parechovirus* | *Parechovirus A* | nonsevere |  | 1 | 0 | 1 | [2,83–85] |
| *Picornaviridae* | *Parechovirus* | *Parechovirus B* | severe | Associated with sudden infant death syndrome and adult myocarditis | 0 | 0 | N/A | [86–89] |
| *Picornaviridae* | *Salivirus* | *Salivirus A* | nonsevere |  | 0 | 0 | 0 | [90–92] |
| *Togaviridae* | *Alphavirus* | *Barmah Forest virus* | nonsevere |  | 0 | 0 | 0 | [4] |
| *Togaviridae* | *Alphavirus* | *Chikungunya virus* | nonsevere |  | 1 | 1 | 0 | [2,4] |
| *Togaviridae* | *Alphavirus* | *Eastern equine encephalitis virus* | severe | CFR 30-70% | 1 | 1 | N/A | [2,4] |
| *Togaviridae* | *Alphavirus* | *Everglades virus* | nonsevere |  | 0 | 0 | 0 | [1,93] |
| *Togaviridae* | *Alphavirus* | *Getah virus* | nonsevere |  | 0 | 0 | 0 | [1,4] |
| *Togaviridae* | *Alphavirus* | *Mayaro virus* | nonsevere |  | 0 | 0 | 0 | [2] |
| *Togaviridae* | *Alphavirus* | *Mosso das Pedras virus* | nonsevere |  | 0 | 0 | 0 | [94] |
| *Togaviridae* | *Alphavirus* | *Ndumu virus* | nonsevere |  | 0 | 0 | 0 | [95] |
| *Togaviridae* | *Alphavirus* | *O'nyong-nyong virus* | nonsevere |  | 0 | 0 | 0 | [1,4,96] |
| *Togaviridae* | *Alphavirus* | *Pixuna virus* | nonsevere |  | 0 | 0 | 0 | [97] |
| *Togaviridae* | *Alphavirus* | *Rio Negro virus* | nonsevere |  | 0 | 0 | 0 | [1,4] |
| *Togaviridae* | *Alphavirus* | *Ross River virus* | nonsevere |  | 0 | 0 | 0 | [1,2,4] |
| *Togaviridae* | *Alphavirus* | *Semliki Forest virus* | nonsevere |  | 0 | 0 | 0 | [1,4] |
| *Togaviridae* | *Alphavirus* | *Tonate virus* | nonsevere |  | 1 | 0 | 0 | [1,4,98,99] |
| *Togaviridae* | *Alphavirus* | *Una virus* | nonsevere |  | 0 | 0 | 0 | [2,4] |
| *Togaviridae* | *Alphavirus* | *Western equine encephalitis virus* | severe | CFR 3-10% | 1 | 1 | N/A | [1,4] |
| *Togaviridae* | *Alphavirus* | *Whataroa virus* | nonsevere |  | 0 | 0 | 0 | [100] |
| *Togaviridae* | *Rubivirus* | *Rubella virus* | nonsevere |  | 1 | 1 | 0 | [1,4] |
| *dsRNA viruses* | |  |  |  |  |  |  |  |
| *Picobirnaviridae* | *Picobirnavirus* | *Human picobirnavirus* | nonsevere |  | 0 | 0 | 0 | [2] |
| *Reoviridae* | *Coltivirus* | *Colorado tick fever virus* | nonsevere |  | 1 | 0 | 0 | [1] |
| *Reoviridae* | *Coltivirus* | *Eyach virus* | nonsevere |  | 0 | 0 | 0 | [101] |
| *Reoviridae* | *Orbivirus* | *Corriparta virus* | nonsevere |  | 0 | 0 | 0 | [102] |
| *Reoviridae* | *Orbivirus* | *Great Island virus* | nonsevere |  | 0 | 0 | 0 | [1,103] |
| *Reoviridae* | *Orbivirus* | *Lebombo virus* | nonsevere |  | 0 | 0 | 0 | [1] |
| *Reoviridae* | *Orthoreovirus* | *Mammalian orthoreovirus* | nonsevere |  | 0 | 0 | 0 | [4] |
| *Reoviridae* | *Orthoreovirus* | *Nelson Bay orthoreovirus* | nonsevere |  | 0 | 0 | 0 | [104–106] |
| *Reoviridae* | *Rotavirus* | *Rotavirus A* | nonsevere |  | 1 | 0 | 0 | [1,4] |
| *Reoviridae* | *Rotavirus* | *Rotavirus B* | nonsevere |  | 1 | 0 | 0 | [4] |
| *Reoviridae* | *Rotavirus* | *Rotavirus H* | nonsevere |  | 0 | 0 | 0 | [107,108] |
| *Reoviridae* | *Seadornavirus* | *Banna virus* | nonsevere |  | 0 | 0 | 0 | [1,109,110] |
| *ssRNA-RT viruses* | |  |  |  |  |  |  |  |
| *Retroviridae* | *Deltaretrovirus* | *Primate T-lymphotropic virus 1* | severe | Chronic neoplastic and neurologic disease including lymphoma, leukaemia, and HTLV-associated myelopathy | 1 | 1 | N/A | [1,4] |
| *Retroviridae* | *Deltaretrovirus* | *Primate T-lymphotropic virus 2* | severe | Chronic neurologic disease including HTLV-associated myelopathy | 1 | 1 | N/A | [1,111] |
| *Retroviridae* | *Lentivirus* | *Human immunodeficiency virus 1* | severe | AIDS | 1 | 1 | N/A | [4] |
| *Retroviridae* | *Lentivirus* | *Human immunodeficiency virus 2* | severe | AIDS | 1 | 1 | N/A | [4] |
| *Retroviridae* | *Lentivirus* | *Simian immunodeficiency virus* | nonsevere |  | 0 | 0 | 0 | [112] |
| *Retroviridae* | *Spumavirus* | *African green monkey simian foamy virus* | nonsevere |  | 0 | 0 | 0 | [4,113] |
| *Retroviridae* | *Spumavirus* | *Macaque simian foamy virus* | nonsevere |  | 0 | 0 | 0 | [4,114,115] |
| *Retroviridae* | *Spumavirus* | *Simian foamy virus* | nonsevere |  | 0 | 0 | 0 | [4] |

References from Table S1:

1.	Richman DD, Whitley RJ, Hayden FG. Clinical virology. John Wiley & Sons; 2009.

2.	Zuckerman AJ, Banatvala JE, Griffiths P, Schoub B, Mortimer P. Principles and practice of clinical virology. John Wiley & Sons; 2009.

3.	Delgado S, Erickson BR, Agudo R, Blair PJ, Vallejo E, Albariño CG, et al. Chapare virus, a newly discovered arenavirus isolated from a fatal hemorrhagic fever case in Bolivia. PLoS Pathog. 2008;4: e1000047.

4.	Knipe DM, Howley PM. Fields virology, 5th Edition. Lippincott Williams & Wilkins; 2007.

5.	Briese T, Paweska JT, McMullan LK, Hutchison SK, Street C, Palacios G, et al. Genetic detection and characterization of Lujo virus, a new hemorrhagic fever–associated arenavirus from southern Africa. PLoS Pathog. 2009;5: e1000455. doi:10.1371/journal.ppat.1000455

6.	Paweska JT, Sewlall NH, Ksiazek TG, Blumberg LH, Hale MJ, Lipkin WI, et al. Nosocomial outbreak of novel arenavirus infection, Southern Africa. Emerg Infect Dis. 2009;15: 1598–1602. doi:10.3201/eid1510.090211

7.	Georges AJ, Gonzalez JP, Abdul-Wahid S, Saluzzo JF, Meunier DMY, McCormick JB. Antibodies to Lassa and lassa-like viruses in man and mammals in the Central African Republic. Trans R Soc Trop Med Hyg. 1985;79: 78–79. doi:10.1016/0035-9203(85)90242-1

8.	Hoffmann B, Tappe D, Höper D, Herden C, Boldt A, Mawrin C, et al. A Variegated Squirrel Bornavirus Associated with Fatal Human Encephalitis. N Engl J Med. 2015;373: 154–162. doi:10.1056/NEJMoa1415627

9.	Towner JS, Sealy TK, Khristova ML, Albariño CG, Conlan S, Reeder SA, et al. Newly discovered ebola virus associated with hemorrhagic fever outbreak in Uganda. PLoS Pathog. 2008;4: e1000212. doi:10.1371/journal.ppat.1000212

10.	MacNeil A, Farnon EC, Wamala J, Okware S, Cannon DL, Reed Z, et al. Proportion of deaths and clinical features in Bundibugyo ebola virus infection, Uganda. Emerg Infect Dis. 2010;16: 1969–1972. doi:10.3201/eid1612.100627

11.	Khan AS, Spiropoulou CF, Morzunov S, Zaki SR, Kohn MA, Nawas SR, et al. Fatal illness associated with a new hantavirus in Louisiana. J Med Virol. 1995;46: 281–286.

12.	Klempa B, Witkowski PT, Popugaeva E, Auste B, Koivogui L, Fichet-Calvet E, et al. Sangassou Virus, the First Hantavirus Isolate from Africa, Displays Genetic and Functional Properties Distinct from Those of Other Murinae-Associated Hantaviruses. J Virol. 2012;86: 3819–3827. doi:10.1128/JVI.05879-11

13.	Pattamadilok S, Lee B-H, Kumperasart S, Yoshimatsu K, Okumura M, Nakamura I, et al. Geographical distribution of hantaviruses in Thailand and potential human health significance of Thailand virus. Am J Trop Med Hyg. 2006;75: 994–1002.

14.	Okumura M, Yoshimatsu K, Kumperasart S, Nakamura I, Ogino M, Taruishi M, et al. Development of Serological Assays for Thottapalayam Virus, an Insectivore-Borne Hantavirus. Clin Vaccine Immunol. 2007;14: 173–181. doi:10.1128/CVI.00347-06

15.	Klempa B, Meisel H, Räth S, Bartel J, Ulrich R, Krüger DH. Occurrence of renal and pulmonary syndrome in a region of northeast Germany where Tula hantavirus circulates. J Clin Microbiol. 2003;41: 4894–4897. doi:10.1128/JCM.41.10.4894-4897.2003

16.	Treib J, Dobler G, Haass A, von Blohn W, Strittmatter M, Pindur G, et al. Thunderclap headache caused by Erve virus? Neurology. 1998;50: 509–511.

17.	Woessner R, Grauer MT, Langenbach J, Dobler G, Kroeger J, Mielke HG, et al. The Erve virus: possible mode of transmission and reservoir. Infection. 2000;28: 164–166.

18.	Butenko AM, Leshchinskaia EV, Semashko IV, Donets MA, Mart’ianova LI. [Dhori virus--a causative agent of human disease. 5 cases of laboratory infection]. Vopr Virusol. 1987;32: 724–729.

19.	Moore DL, Causey OR, Carey DE, Reddy S, Cooke AR, Akinkugbe FM, et al. Arthropod-borne viral infections of man in Nigeria, 1964-1970. Ann Trop Med Parasitol. 1975;69: 49.

20.	Goebel SJ, Taylor J, Barr BC, Kiehn TE, Castro-Malaspina HR, Hedvat CV, et al. Isolation of Avian paramyxovirus 1 from a patient with a lethal case of pneumonia. J Virol. 2007;81: 12709–12714. doi:10.1128/JVI.01406-07

21.	ProMED-mail | Hendra virus, human, equine - Australia (04): (QL) fatal [Internet]. [cited 3 Jul 2014]. Available: http://www.promedmail.org/direct.php?id=283896

22.	Selby PL, Davies M, Mee AP. Canine distemper virus induces human osteoclastogenesis through NF-kappaB and sequestosome 1/P62 activation. J Bone Miner Res Off J Am Soc Bone Miner Res. 2006;21: 1750–1756. doi:10.1359/jbmr.060805

23.	Chatziandreou N, Stock N, Young D, Andrejeva J, Hagmaier K, McGeoch DJ, et al. Relationships and host range of human, canine, simian and porcine isolates of simian virus 5 (parainfluenza virus 5). J Gen Virol. 2004;85: 3007–3016. doi:10.1099/vir.0.80200-0

24.	Albarino CG, Foltzer M, Towner JS, Rowe LA, Campbell S, Jaramillo CM, et al. Novel paramyxovirus associated with severe acute febrile disease, South Sudan and Uganda, 2012. Emerg Infect Dis. 2014;20: 211–216. doi:10.3201/eid2002.131620

25.	Mackenzie JS, Field HE, Guyatt KJ. Managing emerging diseases borne by fruit bats (flying foxes), with particular reference to henipaviruses and Australian bat lyssavirus. J Appl Microbiol. 2003;94: 59–69. doi:10.1046/j.1365-2672.94.s1.7.x

26.	Rodaniche E de, Andrade AP, Galindo P. Isolation of two antigenically distinct arthropod-borne viruses of group C in Panama. Am J Trop Med Hyg. 1964;13: 839–843.

27.	Digoutte JP, Gagnard VJM, Brès P, Pajot F-X. Infection à virus Nyando chez l’homme. Bull Société Pathol Exot. 1972;65: 751–758.

28.	Scherer WF, Anderson K, Dickerman RW, Ordonez JV. Studies of Patois Group Arboviruses in Mexico, Guatemala, Honduras, and British Honduras. Am J Trop Med Hyg. 1972;21: 194–200.

29.	Vasconcelos PFC, Rosa A, Rosa J, Dégallier N. Concomitant infections by malaria and arboviruses in the Brazilian Amazon region. Rev Latinoam Microbiol. 1990;32: 291–294.

30.	Srihongse S, Johnson CM. Wyeomyia subgroup of arbovirus: isolation from man. Science. 1965;149: 863–864.

31.	Palacios G, Tesh R, Travassos da Rosa A, Savji N, Sze W, Jain K, et al. Characterization of the Candiru antigenic complex (Bunyaviridae: Phlebovirus), a highly diverse and reassorting group of viruses affecting humans in tropical America. J Virol. 2011;85: 3811–3820. doi:10.1128/JVI.02275-10

32.	Yu X-J, Liang M-F, Zhang S-Y, Liu Y, Li J-D, Sun Y-L, et al. Fever with thrombocytopenia associated with a novel bunyavirus in China. N Engl J Med. 2011;364: 1523–1532. doi:10.1056/NEJMoa1010095

33.	Xie Q, Li X, Cheng J, Shao Y. Multiple organ damage caused by a novel tick-borne bunyavirus: A case report. J Vector Borne Dis. 2013;50: 314–317.

34.	Zhang Y-Z, Zhou D-J, Xiong Y, Chen X-P, He Y-W, Sun Q, et al. Hemorrhagic fever caused by a novel tick-borne Bunyavirus in Huaiyangshan, China. Zhonghua Liu Xing Bing Xue Za Zhi Zhonghua Liuxingbingxue Zazhi. 2011;32: 209–220.

35.	Zhao L, Zhai S, Wen H, Cui F, Chi Y, Wang L, et al. Severe fever with thrombocytopenia syndrome virus, Shandong Province, China. Emerg Infect Dis. 2012;18: 963–965. doi:10.3201/eid1806.111345

36.	Kayali G, Ortiz EJ, Chorazy ML, Nagaraja KV, DeBeauchamp J, Webby RJ, et al. Serologic evidence of avian metapneumovirus infection among adults occupationally exposed to Turkeys. Vector Borne Zoonotic Dis Larchmt N. 2011;11: 1453–1458. doi:10.1089/vbz.2011.0637

37.	Familusi JB, Osunkoya BO, Moore DL, Kemp GE, Fabiyi A. A fatal human infection with Mokola virus. Am J Trop Med Hyg. 1972;21: 959–963.

38.	Grard G, Fair JN, Lee D, Slikas E, Steffen I, Muyembe J-J, et al. A novel rhabdovirus associated with acute hemorrhagic fever in Central Africa. PLoS Pathog. 2012;8: e1002924. doi:10.1371/journal.ppat.1002924

39.	Stremlau MH, Andersen KG, Folarin OA, Grove JN, Odia I, Ehiane PE, et al. Discovery of novel rhabdoviruses in the blood of healthy individuals from West Africa. PLoS Negl Trop Dis. 2015;9: e0003631. doi:10.1371/journal.pntd.0003631

40.	Letchworth GJ, Rodriguez LL, Del Cbarrera J. Vesicular stomatitis. Vet J. 1999;157: 239–260.

41.	Jonkers AH, Shope RE, Aitken TH, Spence L. COCAL VIRUS, A NEW AGENT IN TRINIDAD RELATED TO VESICULAR STOMATITIS VIRUS, TYPE INDIANA. Am J Vet Res. 1964;25: 236–242.

42.	Travassos da Rosa APA, Tesh RB, Travassos da Rosa JF, Herve JP, Main AJJ. Carajas and Maraba viruses, two new vesiculoviruses isolated from phlebotomine sand flies in Brazil. Am J Trop Med Hyg. 1984;33: 999–1006.

43.	Pedrosa PBS, Cardoso TAO. Viral infections in workers in hospital and research laboratory settings: a comparative review of infection modes and respective biosafety aspects. Int J Infect Dis. 2011;15: e366–e376. doi:10.1016/j.ijid.2011.03.005

44.	Singh PB, Sreenivasan MA, Pavri KM. Viruses in acute gastroenteritis in children in Pune, India. Epidemiol Infect. 1989;102: 345–353.

45.	Guix S, Bosch A, Pintó RM. Astrovirus Taxonomy. In: Schultz-Cherry S, editor. Astrovirus Research. Springer New York; 2013. pp. 97–118.

46.	Finkbeiner SR, Holtz LR, Jiang Y, Rajendran P, Franz CJ, Zhao G, et al. Human stool contains a previously unrecognized diversity of novel astroviruses. Virol J. 2009;6: 161. doi:10.1186/1743-422X-6-161

47.	Ahmed SF, Sebeny PJ, Klena JD, Pimentel G, Mansour A, Naguib AM, et al. Novel astroviruses in children, Egypt. Emerg Infect Dis. 2011;17: 2391–2393. doi:10.3201/eid1712.110909

48.	Phan TG, Vo NP, Bonkoungou IJO, Kapoor A, Barro N, O’Ryan M, et al. Acute diarrhea in West African children: diverse enteric viruses and a novel parvovirus genus. J Virol. 2012;86: 11024–11030. doi:10.1128/JVI.01427-12

49.	Finkbeiner SR, Li Y, Ruone S, Conrardy C, Gregoricus N, Toney D, et al. Identification of a novel astrovirus (Astrovirus VA1) associated with an outbreak of acute gastroenteritis. J Virol. 2009;83: 10836–10839. doi:10.1128/JVI.00998-09

50.	Smith AW, Berry ES, Skilling DE, Barlough JE, Poet SE, Berke T, et al. In vitro isolation and characterization of a calicivirus causing a vesicular disease of the hands and feet. Clin Infect Dis Off Publ Infect Dis Soc Am. 1998;26: 434–439.

51.	Terao Y, Takagi H, Phan TG, Okitsu S, Ushijima H. Identification of antibody against porcine coronavirus in human milk. Clin Lab. 2007;53: 129–130.

52.	Pene F, Merlat A, Vabret A, Rozenberg F, Buzyn A, Dreyfus F, et al. Coronavirus 229E-Related Pneumonia in Immunocompromised Patients. Clin Infect Dis. 2003;37: 929–932. doi:10.1086/377612

53.	Bastien N, Anderson K, Hart L, Caeseele PV, Brandt K, Milley D, et al. Human coronavirus NL63 infection in Canada. J Infect Dis. 2005;191: 503–506. doi:10.1086/426869

54.	Patrick DM, Petric M, Skowronski DM, Guasparini R, Booth TF, Krajden M, et al. An outbreak of Human coronavirus OC43 infection and serological cross-reactivity with SARS coronavirus. Can J Infect Dis Med Microbiol. 2006;17: 330–336.

55.	Zaki AM, Van Boheemen S, Bestebroer TM, Osterhaus AD, Fouchier RA. Isolation of a novel coronavirus from a man with pneumonia in Saudi Arabia. N Engl J Med. 2012;367: 1814–1820.

56.	WHO. Frequently asked questions on Middle East respiratory syndrome coronavirus (MERS-CoV) [Internet]. [cited 10 Sep 2014]. Available: http://www.who.int/csr/disease/coronavirus_infections/faq/en/

57.	Chan JF-W, Lau SK-P, Woo PC-Y. The emerging novel Middle East respiratory syndrome coronavirus: the “knowns” and “unknowns.” J Formos Med Assoc Taiwan Yi Zhi. 2013;112: 372–381. doi:10.1016/j.jfma.2013.05.010

58.	Srihongse S, Johnson CM. The first isolation of Bussuquara virus from man. Trans R Soc Trop Med Hyg. 1971;65: 541–542.

59.	Calisher CH, Gould EA. Taxonomy of the virus family Flaviviridae. Advances in Virus Research. Academic Press; 2003. pp. 1–19.

60.	Batista WC, Tavares G da SB, Vieira DS, Honda ER, Pereira SS, Tada MS. Notification of the first isolation of Cacipacore virus in a human in the State of Rondônia, Brazil. Rev Soc Bras Med Trop. 2011;44: 528–530.

61.	Aaskov JG, Phillips DA, Wiemers MA. Possible clinical infection with Edge Hill virus. Trans R Soc Trop Med Hyg. 1993;87: 452–453.

62.	Růžek D, Yakimenko VV, Karan LS, Tkachev SE. Omsk haemorrhagic fever. The Lancet. 2010;376: 2104–2113. doi:10.1016/S0140-6736(10)61120-8

63.	Weissenböck H, Hubálek Z, Bakonyi T, Nowotny N. Zoonotic mosquito-borne flaviviruses: worldwide presence of agents with proven pathogenicity and potential candidates of future emerging diseases. Vet Microbiol. 2010;140: 271–280.

64.	Bhattarai N, Stapleton JT. GB virus C: the good boy virus? Trends Microbiol. 2012;20: 124–130. doi:10.1016/j.tim.2012.01.004

65.	Giangaspero M, Wellemans G, Vanopdenbosch E, Belloli A, Verhulst A. Bovine viral diarrhoea. The Lancet. 1988;332: 110. doi:10.1016/S0140-6736(88)90047-5

66.	Plummer G. An equine respiratory virus with enterovirus properties. Nature. 1962;195: 519–520. doi:10.1038/195519a0

67.	Bauer K. Foot- and-mouth disease as zoonosis. Arch Virol Suppl. 1997;13: 95–97.

68.	Oberste MS, Gotuzzo E, Blair P, Nix WA, Ksiazek TG, Comer JA, et al. Human febrile illness caused by Encephalomyocarditis virus infection, Peru. Emerg Infect Dis. 2009;15: 640–646. doi:10.3201/eid1504.081428

69.	Lipton HL. Human Vilyuisk encephalitis. Rev Med Virol. 2008;18: 347–352. doi:10.1002/rmv.585

70.	Goldfarb LG, Vladimirtsev VA, Platonov FA, Lee H-S, McLean CA, Masters CL. Viliuisk encephalomyelitis in Eastern Siberia - analysis of 390 cases: In memory of D Carleton Gajdusek. Folia Neuropathol Assoc Pol Neuropathol Med Res Cent Pol Acad Sci. 2009;47: 171–181.

71.	Zoll J, Erkens Hulshof S, Lanke K, Verduyn Lunel F, Melchers WJG, Schoondermark-van de Ven E, et al. Saffold virus, a human Theiler’s-like cardiovirus, is ubiquitous and causes infection early in life. PLoS Pathog. 2009;5: e1000416. doi:10.1371/journal.ppat.1000416

72.	Kapoor A, Li L, Victoria J, Oderinde B, Mason C, Pandey P, et al. Multiple novel astrovirus species in human stool. J Gen Virol. 2009;90: 2965–2972. doi:10.1099/vir.0.014449-0

73.	Kapusinszky B, Phan TG, Kapoor A, Delwart E. Genetic diversity of the genus Cosavirus in the family Picornaviridae: a new species, recombination, and 26 new genotypes. PloS One. 2012;7: e36685. doi:10.1371/journal.pone.0036685

74.	Stocker A, Souza BF de CD, Ribeiro TCM, Netto EM, Araujo LO, Correa JI, et al. Cosavirus infection in persons with and without gastroenteritis, Brazil. Emerg Infect Dis. 2012;18: 656–659. doi:10.3201/eid1804.111415

75.	Dai XQ, Hua XG, Shan TL, Delwart E, Zhao W. Human cosavirus infections in children in China. J Clin Virol. 2010;48: 228–229. doi:10.1016/j.jcv.2010.03.024

76.	Campanini G, Rovida F, Meloni F, Cascina A, Ciccocioppo R, Piralla A, et al. Persistent human cosavirus infection in lung transplant recipient, Italy. Emerg Infect Dis. 2013;19: 1667–1669. doi:10.3201/eid1910.130352

77.	Holtz LR, Finkbeiner SR, Kirkwood CD, Wang D. Identification of a novel picornavirus related to cosaviruses in a child with acute diarrhea. Virol J. 2008;5: 159. doi:10.1186/1743-422X-5-159

78.	Yi L, Lu J, Kung H, He M-L. The virology and developments toward control of human enterovirus 71. Crit Rev Microbiol. 2011;37: 313–327. doi:10.3109/1040841X.2011.580723

79.	Louie JK, Yagi S, Nelson FA, Kiang D, Glaser CA, Rosenberg J, et al. Rhinovirus outbreak in a long term care facility for elderly persons associated with unusually high mortality. Clin Infect Dis Off Publ Infect Dis Soc Am. 2005;41: 262–265. doi:10.1086/430915

80.	Kriegshäuser G, Deutz A, Kuechler E, Skern T, Lussy H, Nowotny N. Prevalence of neutralizing antibodies to Equine rhinitis A and B virus in horses and man. Vet Microbiol. 2005;106: 293–296.

81.	Ambert-Balay K, Lorrot M, Bon F, Giraudon H, Kaplon J, Wolfer M, et al. Prevalence and genetic diversity of Aichi virus strains in stool samples from community and hospitalized patients. J Clin Microbiol. 2008;46: 1252–1258. doi:10.1128/JCM.02140-07

82.	Kaikkonen S, Räsänen S, Rämet M, Vesikari T. Aichi virus infection in children with acute gastroenteritis in Finland. Epidemiol Infect. 2010;138: 1166–1171. doi:10.1017/S0950268809991300

83.	Benschop KSM, Schinkel J, Minnaar RP, Pajkrt D, Spanjerberg L, Kraakman HC, et al. Human parechovirus infections in Dutch children and the association between serotype and disease severity. Clin Infect Dis. 2006;42: 204–210. doi:10.1086/498905

84.	Ito M, Yamashita T, Tsuzuki H, Takeda N, Sakae K. Isolation and identification of a novel human parechovirus. J Gen Virol. 2004;85: 391–398.

85.	Abed Y, Boivin G. Human Parechovirus Infections in Canada. Emerg Infect Dis. 2006;12: 969–975. doi:10.3201/eid1206.051675

86.	Blixt M, Sandler S, Bo Niklasson M. D. PD. Ljungan Virus and Diabetes. In: Taylor K, Hyöty H, Toniolo A, Zuckerman AJ, editors. Diabetes and Viruses. Springer New York; 2013. pp. 81–86.

87.	Krous HF, Langlois NE. Ljungan virus: a commentary on its association with fetal and infant morbidity and mortality in animals and humans. Birt Defects Res A Clin Mol Teratol. 2010;88: 947–952. doi:10.1002/bdra.20728

88.	Samsioe A, Papadogiannakis N, Hultman T, Sjöholm Å, Klitz W, Niklasson B. Ljungan virus present in intrauterine fetal death diagnosed by both immunohistochemistry and PCR. Birt Defects Res A Clin Mol Teratol. 2009;85: 227–229. doi:10.1002/bdra.20554

89.	Niklasson B, Samsioe A, Papadogiannakis N, Gustafsson S, Klitz W. Zoonotic Ljungan virus associated with central nervous system malformations in terminated pregnancy. Birt Defects Res A Clin Mol Teratol. 2009;85: 542–545. doi:10.1002/bdra.20568

90.	Greninger AL, Runckel C, Chiu CY, Haggerty T, Parsonnet J, Ganem D, et al. The complete genome of klassevirus—a novel picornavirus in pediatric stool. Virol J. 2009;6: 82.

91.	Li L, Victoria J, Kapoor A, Blinkova O, Wang C, Babrzadeh F, et al. A novel picornavirus associated with gastroenteritis. J Virol. 2009;83: 12002–12006. doi:10.1128/JVI.01241-09

92.	Shan T, Wang C, Cui L, Yu Y, Delwart E, Zhao W, et al. Picornavirus salivirus/klassevirus in children with diarrhea, China. Emerg Infect Dis. 2010;16: 1303–1305. doi:10.3201/eid1608.100087

93.	Ehrenkranz NJ, Ventura AK. Venezuelan equine encephalitis virus infection in man. Annu Rev Med. 1974;25: 9–14. doi:10.1146/annurev.me.25.020174.000301

94.	Pisano MB, Oria G, Beskow G, Aguilar J, Konigheim B, Cacace ML, et al. Venezuelan Equine Encephalitis Viruses (VEEV) in Argentina: Serological Evidence of Human Infection. PLoS Negl Trop Dis. 2013;7: e2551. doi:10.1371/journal.pntd.0002551

95.	Kokernot RH, McIntosh BM, Worth CB. Ndumu virus, a hitherto unknown agent, isolated from culicine mosouitoes collected in northern Natal. Union of South Africa. Am J Trop Med Hyg. 1961;10: 383–386.

96.	Kiwanuka N, Sanders EJ, Rwaguma EB, Kawamata J, Ssengooba FP, Najjemba R, et al. O’nyong-nyong fever in south-central Uganda, 1996—1997: clinical features and validation of a clinical case definition for surveillance purposes. Clin Infect Dis. 1999;29: 1243–1250. doi:10.1086/313462

97.	Vasconcelos PF da C, Travassos da Rosa JFS, Travassos da Rosa AP de A, Dégallier N, Pinheiro F de P, Sá Filho GC. Epidemiology of encephalitis by arboviruses in the Amazon region of Brazil. Rev Inst Med Trop São Paulo. 1991;33: 465–476. doi:10.1590/S0036-46651991000600007

98.	Hommel D, Heraud JM, Hulin A, Talarmin A. Association of Tonate virus (subtype IIIB of the Venezuelan equine encephalitis complex) with encephalitis in a human. Clin Infect Dis. 2000;30: 188–190. doi:10.1086/313611

99.	Talarmin A, Trochu J, Gardon J, Laventure S, Hommel D, Lelarge J, et al. Tonate virus infection in French Guiana: clinical aspects and seroepidemiologic study. Am J Trop Med Hyg. 2001;64: 274–279.

100.	Maguire T, Miles JAR, Casals J. Whataroa virus, a group A arbovirus isolated in South Westland, New Zealand. Am J Trop Med Hyg. 1967;16: 371–373.

101.	Dobler G. Arboviruses causing neurological disorders in the central nervous system. Arch Virol Suppl. 1996;11: 33–40.

102.	Boughton CR, Hawkes RA, Naim HM. Arbovirus infection in humans in NSW: seroprevalence and pathogenicity of certain Australian bunyaviruses. Aust N Z J Med. 1990;20: 51–55.

103.	Libíková H, Heinz F, Ujházyová D, Stünzner D. Orbiviruses of the Kemerovo complex and neurological diseases. Med Microbiol Immunol (Berl). 1978;166: 255–263. doi:10.1007/BF02121159

104.	Chua KB, Crameri G, Hyatt A, Yu M, Tompang MR, Rosli J, et al. A previously unknown reovirus of bat origin is associated with an acute respiratory disease in humans. Proc Natl Acad Sci. 2007;104: 11424–11429. doi:10.1073/pnas.0701372104

105.	Chua KB, Voon K, Crameri G, Tan HS, Rosli J, McEachern JA, et al. Identification and characterization of a new orthoreovirus from patients with acute respiratory infections. PLoS ONE. 2008;3: e3803. doi:10.1371/journal.pone.0003803

106.	Chua KB, Voon K, Yu M, Keniscope C, Abdul Rasid K, Wang L-F. Investigation of a potential zoonotic transmission of orthoreovirus associated with acute influenza-like illness in an adult patient. PLoS ONE. 2011;6: e25434. doi:10.1371/journal.pone.0025434

107.	Yang H, Chen S, Ji S. [A novel rotavirus causing large scale of adult diarrhea in Shi Jiazhuang]. Zhonghua Liu Xing Bing Xue Za Zhi Zhonghua Liuxingbingxue Zazhi. 1998;19: 336–338.

108.	Ji S, Bi Y, Yang H, Yang F, Song J, Tao X, et al. [Cultivation and serial propagation of a new rotavirus causing adult diarrhea in primary human embryo kidney cells]. Zhonghua Yi Xue Za Zhi. 2002;82: 14–18.

109.	Liu H, Li MH, Zhai YG, Meng WS, Sun XH, Cao YX, et al. Banna Virus, China, 1987–2007. Emerg Infect Dis. 2010;16: 514.

110.	Liu H, Gao X, Liang G. Newly recognized mosquito-associated viruses in mainland China, in the last two decades. Virol J. 2011;8: 68.

111.	Araujo A, Hall WW. Human T-lymphotropic virus type II and neurological disease. Ann Neurol. 2004;56: 10–19. doi:10.1002/ana.20126

112.	Khabbaz RF, Heneine W, George JR, Parekh B, Rowe T, Woods T, et al. Infection of a Laboratory Worker with Simian Immunodeficiency Virus. N Engl J Med. 1994;330: 172–177. doi:10.1056/NEJM199401203300304

113.	Heneine W, Switzer WM, Sandstrom P, Brown J, Vedapuri S, Schable CA, et al. Identification of a human population infected with simian foamy viruses. Nat Med. 1998;4: 403–407. doi:10.1038/nm0498-403

114.	Brooks JI, Rud EW, Pilon RG, Smith JM, Switzer WM, Sandstrom PA. Cross-species retroviral transmission from macaques to human beings. The Lancet. 2002;360: 387–388. doi:10.1016/S0140-6736(02)09597-1

115.	Jones-Engel L, Engel GA, Schillaci MA, Rompis A, Putra A, Suaryana KG, et al. Primate-to-human retroviral transmission in Asia. Emerg Infect Dis. 2005;11: 1028–1035. doi:10.3201/eid1107.040957
