## Supplemental Table S2 for "Tissue Tropism and Transmission Ecology Predict Virulence of Human RNA Viruses"

| Family | Genus | Species | Severity rating (literature protocol) | Severity notes | Fatalities (vulnerable) | Fatalities (healthy adults) | Severe strains | P(severe) | References |
| --- | --- | --- | --- | --- | --- | --- | --- | --- | --- |
| *-ssRNA viruses* | | |  |  |  |  |  |  |  |
| *Arenaviridae* | *Mammarenavirus* | *Lassa mammarenavirus* | severe | Haemorrhagic fever, CFR 15% | 1 | 1 | N/A | 0.882 | [1] |
| *Arenaviridae* | *Mammarenavirus* | *Whitewater Arroyo mammarenavirus* | severe | Haemorrhagic fever, CFR 100% (3 cases) | 1 | 1 | N/A | 0.863 | [1–3] |
| *Filoviridae* | *Ebolavirus* | *Zaire ebolavirus* | severe | Haemorrhagic fever, CFR 61-90% | 1 | 1 | N/A | 0.792 | [1] |
| *Hantaviridae* | *Orthohantavirus* | *Choclo orthohantavirus* | severe | HPS, CFR 10-25% | 1 | 1 | N/A | 0.973 | [4–6] |
| *Hantaviridae* | *Orthohantavirus* | *Laguna Negra orthohantavirus* | severe | HPS | 1 | 1 | N/A | 0.973 | [1,3,7] |
| *Hantaviridae* | *Orthohantavirus* | *Puumala orthohantavirus* | nonsevere |  | 1 | 1 | 0 | 0.431 | [1,8] |
| *Paramyxoviridae* | *Rubulavirus* | *Simian rubulavirus* | nonsevere |  | 0 | 0 | 0 | 0.064 | [9] |
| *Peribunyaviridae* | *Orthobunyavirus* | *Guaroa orthobunyavirus* | nonsevere |  | 0 | 0 | 0 | 0.094 | [8] |
| *Peribunyaviridae* | *Orthobunyavirus* | *Kairi orthobunyavirus* | nonsevere |  | 0 | 0 | 0 | 0.002 | [10] |
| *Peribunyaviridae* | *Orthobunyavirus* | *Marituba orthobunyavirus* | nonsevere |  | 0 | 0 | 0 | 0.000 | [8,11] |
| *Peribunyaviridae* | *Orthobunyavirus* | *Oriboca orthobunyavirus* | nonsevere |  | 0 | 0 | 0 | 0.000 | [8,11] |
| *Phenuiviridae* | *Phlebovirus* | *Uukuniemi phlebovirus* | nonsevere |  | 0 | 0 | 0 | 0.190 | [8] |
| *Rhabdoviridae* | *Lyssavirus* | *Irkut lyssavirus* | severe | Rabies-like disease, CFR 100% (1 case) | 1 | 1 | N/A | 0.956 | [12] |
| *Rhabdoviridae* | *Vesiculovirus* | *Chandipura vesiculovirus* | nonsevere |  | 1 | 0 | 0 | 0.455 | [1] |
| *Rhabdoviridae* | *Vesiculovirus* | *Isfahan vesiculovirus* | nonsevere |  | 0 | 0 | 0 | 0.010 | [3] |
| *+ssRNA viruses* | |  |  |  |  |  |  |  |  |
| *Astroviridae* | *Mamastrovirus* | *Mamastrovirus 6* | nonsevere |  | 0 | 0 | 0 | 0.009 | [13–15] |
| *Flaviviridae* | *Flavivirus* | *Langat virus* | nonsevere |  | 0 | 0 | 0 | 0.012 | [1,3,8,16] |
| *Flaviviridae* | *Flavivirus* | *Ntaya virus* | nonsevere |  | 0 | 0 | 0 | 0.169 | [17] |
| *Flaviviridae* | ***Flavivirus*** | ***Rio Bravo virus*** | **severe** | Two of the five known cases required hospitalisation | 0 | 0 | N/A | **0.260** | [1,18] |
| *Flaviviridae* | *Flavivirus* | *Tembusu virus* | nonsevere |  | 0 | 0 | 0 | 0.102 | [19] |
| *Flaviviridae* | ***Flavivirus*** | ***Usutu virus*** | **nonsevere** |  | 0 | 0 | 0 | **0.672** | [1,3,20,21] |
| *Flaviviridae* | ***Flavivirus*** | ***Yellow fever virus*** | **severe** | CFR 11-50%, haemorrhagic fever | 1 | 1 | N/A | **0.320** | [1,3,8] |
| *Picornaviridae* | *Cosavirus* | *Cosavirus D* | nonsevere |  | 0 | 0 | 0 | 0.000 | [22–26] |
| *Picornaviridae* | *Enterovirus* | *Enterovirus D* | nonsevere |  | 1 | 0 | 1 | 0.036 | [1,27] |
| *Picornaviridae* | *Enterovirus* | *Enterovirus E* | nonsevere |  | 0 | 0 | 0 | 0.032 | [28] |
| *Togaviridae* | *Alphavirus* | *Highlands J virus* | nonsevere |  | 0 | 0 | 0 | 0.051 | [29] |
| *Togaviridae* | *Alphavirus* | *Mucambo virus* | nonsevere |  | 0 | 0 | 0 | 0.006 | [8,30] |
| *Togaviridae* | *Alphavirus* | *Sindbis virus* | nonsevere |  | 0 | 0 | 0 | 0.086 | [1,3,8] |
| *Togaviridae* | *Alphavirus* | *Venezuelan equine encephalitis virus* | nonsevere |  | 1 | 1 | 1 | 0.427 | [1,3,8,31] |
| *dsRNA viruses* |  |  |  |  |  |  |  |  |  |
| *Reoviridae* | *Orbivirus* | *Orungo virus* | nonsevere |  | 0 | 0 | 0 | 0.024 | [3] |
| *Reoviridae* | *Rotavirus* | *Rotavirus C* | nonsevere |  | 0 | 0 | 0 | 0.013 | [1,3] |

References from Table S2:

1.	Knipe DM, Howley PM. Fields virology, 5th Edition. Lippincott Williams & Wilkins; 2007.

2.	CDC. Fatal illnesses associated with a new world arenavirus--California, 1999-2000. MMWR Morb Mortal Wkly Rep. 2000;49: 709–711.

3.	Richman DD, Whitley RJ, Hayden FG. Clinical virology. John Wiley & Sons; 2009.

4.	Vincent MJ, Quiroz E, Gracia F, Sanchez AJ, Ksiazek TG, Kitsutani PT, et al. Hantavirus pulmonary syndrome in Panama: identification of novel hantaviruses and their likely reservoirs. Virology. 2000;277: 14–19. doi:10.1006/viro.2000.0563

5.	Gracia F, Armien B, Simpson SQ, Munoz C, Broce C, Pascale JM, et al. Convalescent pulmonary dysfunction following hantavirus pulmonary syndrome in Panama and the United States. Lung. 2010;188: 387–391. doi:10.1007/s00408-010-9245-4

6.	Nelson R, Cañate R, Pascale JM, Dragoo JW, Armien B, Armien AG, et al. Confirmation of Choclo virus as the cause of hantavirus cardiopulmonary syndrome and high serum antibody prevalence in Panama. J Med Virol. 2010;82: 1586–1593. doi:10.1002/jmv.21864

7.	Johnson AM, Bowen MD, Ksiazek TG, Williams RJ, Bryan RT, Mills JN, et al. Laguna Negra virus associated with HPS in western Paraguay and Bolivia. Virology. 1997;238: 115–127.

8.	Zuckerman AJ, Banatvala JE, Griffiths P, Schoub B, Mortimer P. Principles and practice of clinical virology. John Wiley & Sons; 2009.

9.	Itoh H, Morimoto Y, Doi Y, Sanpe T. Studies on simian viruses. Some properties of SV41 grown in Vero cell cultures and search for serum neutralizing antibodies in humans and various animals. Virus. 1968;18: 495–503.

10.	Tauro LB, Almeida FL, Contigiani MS. First detection of human infection by Cache Valley and Kairi viruses (Orthobunyavirus) in Argentina. Trans R Soc Trop Med Hyg. 2009;103: 197–199.

11.	Causey OR, Causey CE, Maroja OM, Macedo DG. The isolation of arthropod-borne viruses, including members of two hitherto undescribed serological groups, in the Amazon region of Brazil. Am J Trop Med Hyg. 1961;10: 227–249.

12.	Leonova GN, Belikov SI, Kondratov IG, Krylova NV, Pavlenko EV, Romanova EV, et al. A fatal case of bat lyssavirus infection in Primorye Territory of the Russian Far East. Rabies Bull Eur. 2009;33: 5–8.

13.	Wylie KM, Mihindukulasuriya KA, Sodergren E, Weinstock GM, Storch GA. Sequence analysis of the human virome in febrile and afebrile children. PLoS ONE. 2012;7: e27735. doi:10.1371/journal.pone.0027735

14.	Guix S, Bosch A, Pintó RM. Astrovirus Taxonomy. In: Schultz-Cherry S, editor. Astrovirus Research. Springer New York; 2013. pp. 97–118.

15.	Finkbeiner SR, Holtz LR, Jiang Y, Rajendran P, Franz CJ, Zhao G, et al. Human stool contains a previously unrecognized diversity of novel astroviruses. Virol J. 2009;6: 161. doi:10.1186/1743-422X-6-161

16.	Maffioli C, Grandgirard D, Leib SL, Engler O. SiRNA inhibits replication of Langat virus, a member of the Tick-borne encephalitis virus complex in organotypic rat brain slices. PLoS ONE. 2012;7: e44703. doi:10.1371/journal.pone.0044703

17.	Woodruff AW, Bowen ET, Platt GS. Viral infections in travellers from tropical Africa. Br Med J. 1978;1: 956–958.

18.	Sulkin SE, Burns KF, Shelton DF, Wallis C. Bat salivary gland virus: infections of man and monkey. Tex Rep Biol Med. 1962;20: 113.

19.	Wolfe ND, Kilbourn AM, Karesh WB, Rahman HA, Bosi EJ, Cropp BC, et al. Sylvatic transmission of arboviruses among Bornean orangutans. Am J Trop Med Hyg. 2001;64: 310–316.

20.	Cavrini F, Gaibani P, Longo G, Pierro AM, Rossini G, Bonilauri P, et al. Usutu virus infection in a patient who underwent orthotropic liver transplantation, Italy, August-September 2009. Euro Surveill Bull Eur Sur Mal Transm Eur Commun Dis Bull. 2009;14: 19448.

21.	Pecorari M, Longo G, Gennari W, Grottola A, Sabbatini AM, Tagliazucchi S, et al. First human case of Usutu virus neuroinvasive infection, Italy, August-September 2009. Eurosurveillance. 2009;14: 19446.

22.	Kapoor A, Li L, Victoria J, Oderinde B, Mason C, Pandey P, et al. Multiple novel astrovirus species in human stool. J Gen Virol. 2009;90: 2965–2972. doi:10.1099/vir.0.014449-0

23.	Kapusinszky B, Phan TG, Kapoor A, Delwart E. Genetic diversity of the genus Cosavirus in the family Picornaviridae: a new species, recombination, and 26 new genotypes. PloS One. 2012;7: e36685. doi:10.1371/journal.pone.0036685

24.	Khamrin P, Chaimongkol N, Malasao R, Suantai B, Saikhruang W, Kongsricharoern T, et al. Detection and molecular characterization of cosavirus in adults with diarrhea, Thailand. Virus Genes. 2012;44: 244–246. doi:10.1007/s11262-011-0700-y

25.	Stocker A, Souza BF de CD, Ribeiro TCM, Netto EM, Araujo LO, Correa JI, et al. Cosavirus infection in persons with and without gastroenteritis, Brazil. Emerg Infect Dis. 2012;18: 656–659. doi:10.3201/eid1804.111415

26.	Cotten M, Oude Munnink B, Canuti M, Deijs M, Watson SJ, Kellam P, et al. Full genome virus detection in fecal samples using sensitive nucleic acid preparation, deep sequencing, and a novel iterative sequence classification algorithm. PLoS ONE. 2014;9: e93269. doi:10.1371/journal.pone.0093269

27.	Imamura T. Clusters of acute respiratory illness associated with human enterovirus 68--Asia, Europe, and United States, 2008-2010. MMWR Morb Mortal Wkly Rep. 2011;60: 1301–1304.

28.	Gür S, Yapkiç O, Yilmaz A. Serological survey of bovine enterovirus type 1 in different mammalian species in Turkey. Zoonoses Public Health. 2008;55: 106–111. doi:10.1111/j.1863-2378.2007.01095.x

29.	Meehan PJ, Wells DL, Paul W, Buff E, Lewis A, Muth D, et al. Epidemiological features of and public health response to a St. Louis encephalitis epidemic in Florida, 1990-1. Epidemiol Infect. 2000;125: 181–188. doi:10.2307/3864999

30.	Vasconcelos PF da C, Travassos da Rosa JFS, Travassos da Rosa AP de A, Dégallier N, Pinheiro F de P, Sá Filho GC. Epidemiology of encephalitis by arboviruses in the Amazon region of Brazil. Rev Inst Med Trop São Paulo. 1991;33: 465–476. doi:10.1590/S0036-46651991000600007

31.	Ehrenkranz NJ, Ventura AK. Venezuelan equine encephalitis virus infection in man. Annu Rev Med. 1974;25: 9–14. doi:10.1146/annurev.me.25.020174.000301
