## Supplemental Table S3 for "Tissue Tropism and Transmission Ecology Predict Virulence of Human RNA Viruses"

| Predictor | Trait | Partial dependence | P(severe) |
| --- | --- | --- | --- |
| Family | *Arenaviridae* | -0.322 | 0.366 |
|  | *Bornaviridae* | -0.197 | 0.408 |
|  | *Filoviridae* | -0.199 | 0.403 |
|  | *Hantaviridae* | -0.066 | 0.448 |
|  | *Nairoviridae* | -0.489 | 0.330 |
|  | *Orthomyxoviridae* | -0.754 | 0.240 |
|  | *Paramyxoviridae* | -0.635 | 0.288 |
|  | *Peribunyaviridae* | -0.943 | 0.280 |
|  | *Phenuiviridae* | -0.540 | 0.306 |
|  | *Pneumoviridae* | -0.571 | 0.302 |
|  | *Rhabdoviridae* | -0.472 | 0.359 |
|  | *Astroviridae* | -0.493 | 0.331 |
|  | *Caliciviridae* | -0.591 | 0.301 |
|  | *Coronaviridae* | -0.573 | 0.310 |
|  | *Flaviviridae* | -0.607 | 0.308 |
|  | *Hepeviridae* | -0.276 | 0.381 |
|  | *Picornaviridae* | -0.951 | 0.249 |
|  | *Togaviridae* | -0.965 | 0.248 |
|  | *Picobirnaviridae* | -0.298 | 0.376 |
|  | *Reoviridae* | -1.010 | 0.219 |
|  | *Retroviridae* | -0.195 | 0.408 |
| Genome type | (-)ssRNA | -0.772 | 0.290 |
|  | (+)ssRNA | -1.030 | 0.260 |
|  | dsRNA | -1.031 | 0.245 |
|  | ssRNA-RT | -0.668 | 0.284 |
| Transmissibility level | 2 | -0.951 | 0.276 |
|  | 3 | -0.437 | 0.336 |
|  | 4 | -0.957 | 0.260 |
| Transmission: primary route | direct contact | -0.462 | 0.331 |
|  | faecal-oral | -1.196 | 0.232 |
|  | respiratory | -0.408 | 0.340 |
|  | vector | -1.075 | 0.245 |
| Transmission: direct contact | 0 | -1.069 | 0.269 |
|  | 1 | -0.672 | 0.292 |
| Transmission: faecal-oral | 0 | -0.947 | 0.279 |
|  | 1 | -0.893 | 0.271 |
| Transmission: respiratory | 0 | -1.030 | 0.277 |
|  | 1 | -0.715 | 0.283 |
| Transmission: vector | 0 | -0.648 | 0.310 |
|  | 1 | -1.059 | 0.251 |
| Transmission: multiple routes | 0 | -0.975 | 0.287 |
|  | 1 | -1.055 | 0.256 |
| Transmission: food-borne | 0 | -1.047 | 0.275 |
|  | 1 | -0.787 | 0.286 |
| Transmission: vertical | 0 | -1.039 | 0.273 |
|  | 1 | -0.624 | 0.298 |
| Tropism: primary | circulatory | -0.626 | 0.277 |
|  | gastrointestinal | -1.234 | 0.213 |
|  | hepatic | -0.656 | 0.274 |
|  | neural | -0.134 | 0.424 |
|  | respiratory | -1.084 | 0.216 |
|  | systemic | -0.214 | 0.395 |
|  | vascular | -0.718 | 0.251 |
|  | viraemic | -1.088 | 0.228 |
| Tropism: vascular | 0 | -1.032 | 0.278 |
|  | 1 | -0.776 | 0.278 |
| Tropism: circulatory | 0 | -1.038 | 0.275 |
|  | 1 | -0.724 | 0.286 |
| Tropism: gastrointestinal | 0 | -0.996 | 0.274 |
|  | 1 | -0.772 | 0.291 |
| Tropism: hepatic | 0 | -1.058 | 0.270 |
|  | 1 | -0.464 | 0.326 |
| Tropism: neural | 0 | -1.138 | 0.250 |
|  | 1 | -0.523 | 0.321 |
| Tropism: respiratory | 0 | -1.016 | 0.276 |
|  | 1 | -0.795 | 0.281 |
| Tropism: cardiac | 0 | -1.037 | 0.276 |
|  | 1 | -0.738 | 0.283 |
| Tropism: joints | 0 | -1.036 | 0.276 |
|  | 1 | -0.969 | 0.269 |
| Tropism: renal | 0 | -1.099 | 0.260 |
|  | 1 | -0.176 | 0.404 |
| Tropism: reproductive | 0 | -1.035 | 0.276 |
|  | 1 | -0.959 | 0.270 |
| Tropism: sensory | 0 | -1.037 | 0.276 |
|  | 1 | -0.817 | 0.277 |
| Tropism: skin | 0 | -1.039 | 0.275 |
|  | 1 | -0.664 | 0.291 |
| Tropism: muscular | 0 | -1.050 | 0.274 |
|  | 1 | -0.635 | 0.299 |
| Tropism: endocrine | 0 | -1.055 | 0.271 |
|  | 1 | -0.437 | 0.332 |
| Tropism: multiple tropisms | 0 | -1.038 | 0.275 |
|  | 1 | -0.709 | 0.288 |
| Host range | broad | -0.961 | 0.281 |
|  | narrow | -1.086 | 0.254 |
| Host: human only | 0 | -1.013 | 0.275 |
|  | 1 | -0.929 | 0.275 |
| Host: non-human primates | 0 | -0.958 | 0.285 |
|  | 1 | -1.058 | 0.255 |
| Host: other mammal | 0 | -1.139 | 0.245 |
|  | 1 | -0.930 | 0.284 |
| Host: bird | 0 | -1.135 | 0.260 |
|  | 1 | -0.560 | 0.310 |
