## Supplemental Table S5 for "Tissue Tropism and Transmission Ecology Predict Virulence of Human RNA Viruses"

| Rank | Definition | Example virus species | No. virus species |
| --- | --- | --- | --- |
| 1 | Fits any of ‘severe’ criteria outlined in main text (≥5% case fatality ratio, frequent reports of hospitalisation, significant morbidity from certain symptoms, otherwise explicitly described as “severe” or causing “severe disease”) | *Rabies virus* | 58 |
| 2 | Those not fitting ‘severe’ criteria, but are reported to have caused fatalities in healthy adults | *Dengue virus* | 14 |
| 3 | Those not fitting ‘severe’ criteria, but have severe strains or subspecies reported to cause fatalities in healthy adults | *Influenza A virus* | 6 |
| 4 | Those not fitting ‘severe’ criteria, but are reported to have caused fatalities in vulnerable individuals (age 16 and below or 60 and above, immunosuppressed, having co-morbidities, or otherwise ‘at-risk’) | *Rotavirus A* | 17 |
| 5 | Those not fitting ‘severe’ criteria, but have severe strains or subspecies reported to cause fatalities in vulnerable individuals | *Parechovirus A* | 3 |
| 6 | Those not fitting ‘severe’ criteria that have never been reported to cause fatalities | *Human respirovirus 1* | 114 |
